## Supplementary Materials, Table Legends and Figures for "All small things considered: the diversity of fungi, bacteria and oomycota isolated from the seagrass, *Zostera marina*"

### Supplemental Table and File Legends:

**Table S1:** Media used to isolate microbes associated with *Zostera marina*
Here we report the specifics of the culture media used to initially grow each isolate including the media recipe used, the salt source and amount, and the inclusion of dehydrated crushed seagrass and of various antibiotics.

**Table S2:** Fungal sequences used in molecular phylogenies found based on top BLAST matches to *Zostera marina* associated fungal isolates
Here we report the GenBank accession number and taxonomic information (Class, Order, Molecular ID) for each fungal 28S rRNA gene sequence obtained based on top BLAST matches to fungal isolates in Table 1 and used here to generate Figures 3-6.

**Table S3:** Sequences from fungi isolated from seagrasses used in molecular phylogenies
Here we report information about the fungal 28S rRNA gene sequences used here to generate **Figures 3-6** which represent fungal strains previously isolated from seagrasses. We note the seagrass species and tissue material (e.g. leaf, root, matte or rhizomes) the fungus was isolated from, as well as the taxonomic information (Class, Order, Molecular ID, Strain), GenBank accession number and the study of origin for each fungal 28S rRNA gene sequence.

**Table S4:** Non-seagrass associated fungal isolate sequences from the literature used in molecular phylogenies
Here we report the GenBank accession number and taxonomic information (Phylum, Class, Order, Species, Strain) for each fungal 28S rRNA gene sequence previously used in James et al (2006a,b) and used here to generate Figures 3-6.

**File S1:** R Markdown file of all data analysis performed in R

**Supplemental Figures:**

**Figure S1:** Distribution of counts of fungal isolates across media recipes used for isolation

A histogram representing the number of fungal isolates grouped by order and colored by media recipe used for isolation. The media recipes used included 1% tryptone agar, potato dextrose agar (PDA), potato carrot agar (PCA), palm oil media, lecithin media, malt extract agar (MEA), and glucose yeast peptone agar (GYPA). The numbers included on each bar represent the count of isolates grown on each media recipe.

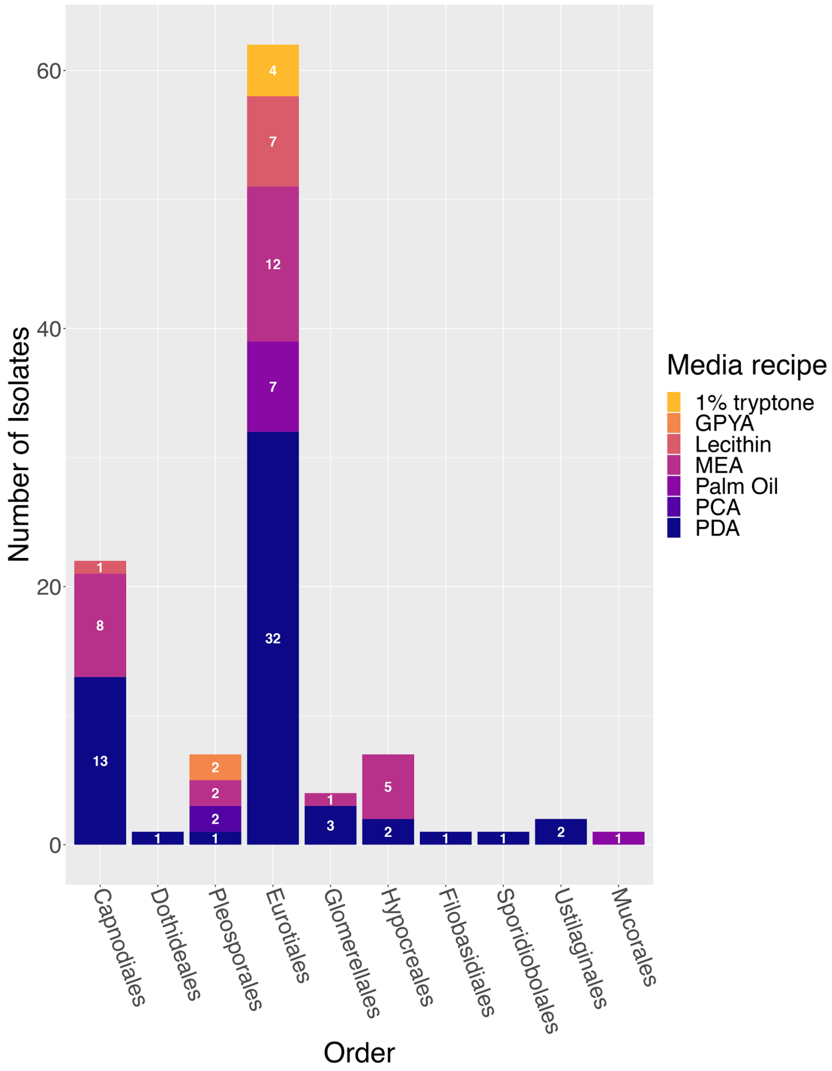

**Figure S2**: Scatterplots showing observed trend between the count of unique media types, salt sources and isolation sources from which a fungal genera was isolated Scatter plots representing A) the relationship between the count of unique isolation sources (leaf, root, rhizome, sediment) and the count of unique media types (PDA, palm oil media, lecithin media, MEA), a fungal genus was isolated from ($R^2$ = 0.86), B) the relationship between the the count of unique isolation sources and the count of unique salt sources (no salt, varying amounts of instant ocean [8 g, 16 g, or 32 g], seawater) a fungal genus was isolated from ($R^2$ = 0.93), and C) the relationship between the count of unique media types and the count of unique salt sources a fungal genus was isolated from ($R^2$ = 0.87).

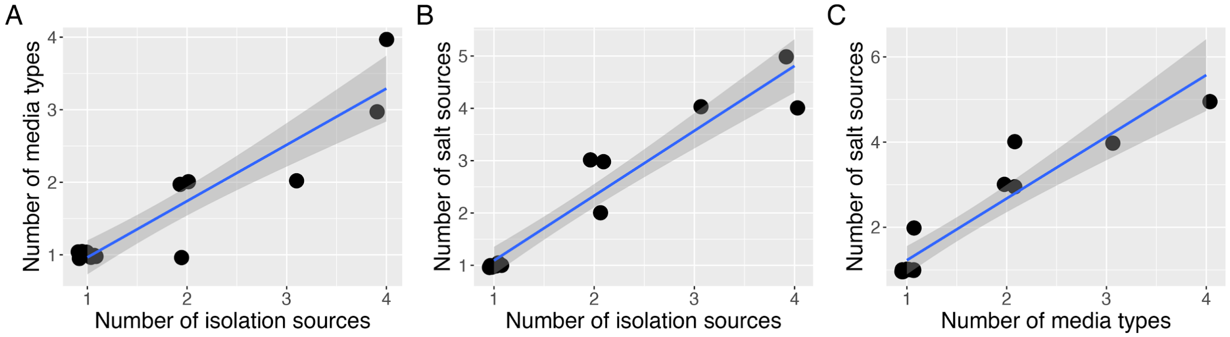

**Figure S3**: Distribution of counts of bacterial isolates across isolation sources and media recipes used for isolation

Histograms representing the number of bacterial isolates grouped by order and colored by isolation source (A) or media recipe (B). A) When colored by isolation source (leaf, leaves and roots, rhizome, root, seawater or sediment), the numbers included on each bar represent the count of isolates obtained from that particular isolation source. B) When colored by media recipe used for isolation (1% tryptone agar, potato dextrose agar [PDA], palm oil media, lecithin media, malt extract agar [MEA], and glucose yeast peptone agar [GYPA]), the numbers included on each bar represent the count of isolates grown on each media recipe.

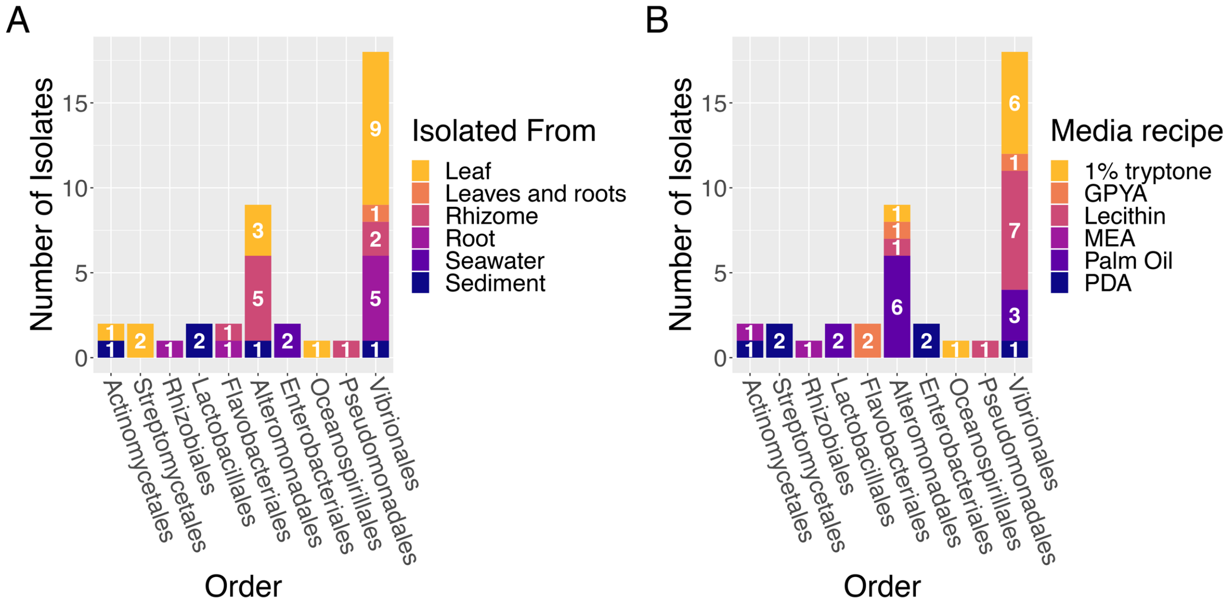

**Figure S4**: Mean relative abundance of fungal orders isolated in this study across sample types in high throughput sequencing data from Ettinger & Eisen (2019)
A histogram representing the mean relative abundance of amplicon sequence variants (ASVs) grouped by order and colored by sample type (leaf, rhizome, root, or sediment). The numbers included on each bar represent the mean relative abundance of the order detected on that particular sample type and only mean relative abundances greater than one percent are shown.

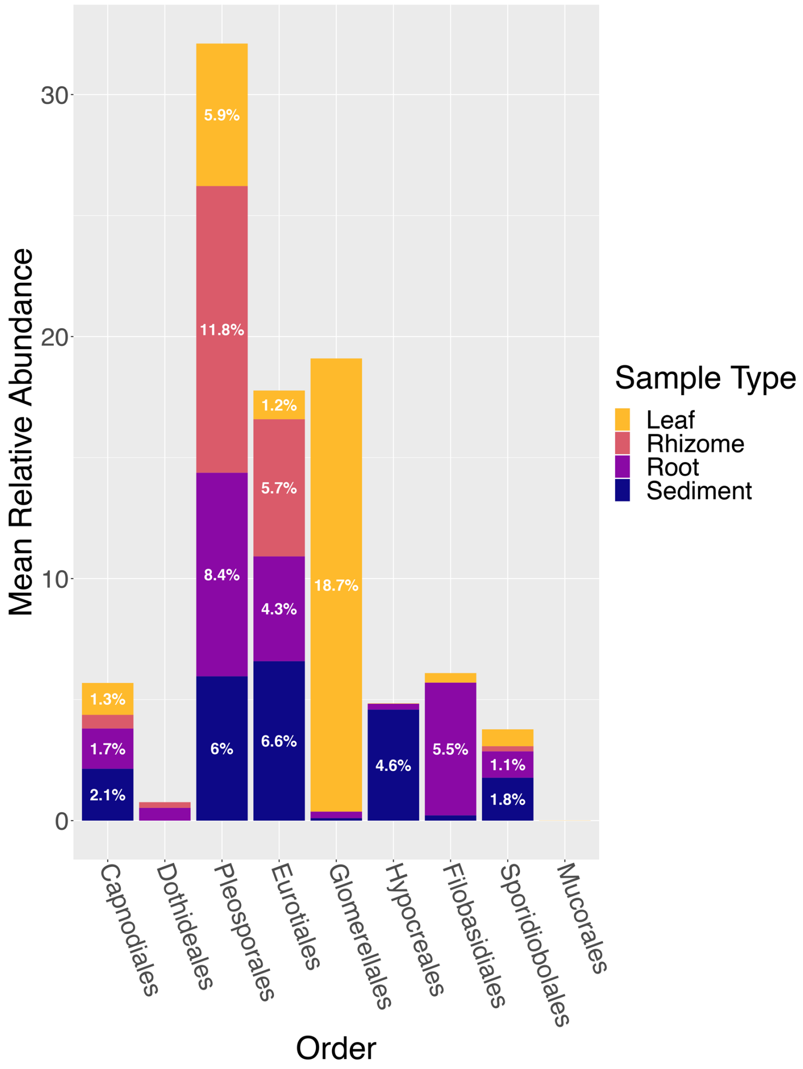
