## Supplemental RMarkdown File for "All small things considered: the diversity of fungi, bacteria and oomycota isolated from the seagrass, *Zostera marina*"

### Seagrass Fungal Isolate Collection R Analysis

Cassie Ettinger

#### Loading packages and setting up the analysis

First, load in the R packages that will be used and make note of their versions.

```
library(tidyverse)
library(ggtree)
library(treeio)
library(dplyr)
library(reshape)
library(ggplot2)
library(patchwork)
library(phyloseq)

# packageVersion('tidyverse') #1.3.0 packageVersion('ggtree')
# #2.0.1 packageVersion('treeio') #1.10.0
# packageVersion('dplyr') #0.8.4 packageVersion('reshape')
# #0.8.8 packageVersion('ggplot2') #3.2.1
# packageVersion('patchwork') #1.0.0
# packageVersion('phyloseq') #1.30.0
```

#### Visualizing the Eurotiomycetes tree

```
# Read in the rooted tree file and the csv containing the
# mapping information (e.g. tip labels, seagrass species,
# tissue isolated from, etc)
euro <- read.mrbayes("euro_v2_rooted.tre")
euro_meta <- read.csv("Euro_data.csv")

# Join metadata with the tree
euro_v2 <- full_join(euro, euro_meta, by = "label")

# Plot the phylogeny and then save as a pdf
p = ggtree(euro_v2, color = "black", size = 1.5, linetype = 1) +
    geom_tiplab(aes(label = Tree_Name2, color = SeagrassREF2),
        fontface = "bold.italic", size = 6, offset = 0.1)
p = p + theme(legend.position = c(0.15, 0.8), legend.text = element_text(size = 24,
    face = "italic"), legend.title = element_text(size = 24)) +
    guides(color = guide_legend(title = "Seagrass Species"))
p = p + xlim(0, 6) + scale_color_manual(values = c(Zostera = "#009E73",
    Reference = "#999999", Seagrass = "#000000")) + geom_point2(aes(subset = !isTip &
    !is.na(as.numeric(prob_percent)), fill = cut(as.numeric(prob_percent),
    c(0, 70, 90, 100))), shape = 21, size = 5) + scale_fill_manual(values = c("white",
    "grey", "black"), guide = "legend", name = "Bayesian Probability (BP)",
```

```
    breaks = c("(90,100]", "(70,90]", "(0,70]"), labels = expression(BP >=
        90, 70 <= BP * " < 90", BP < 70))
p
```

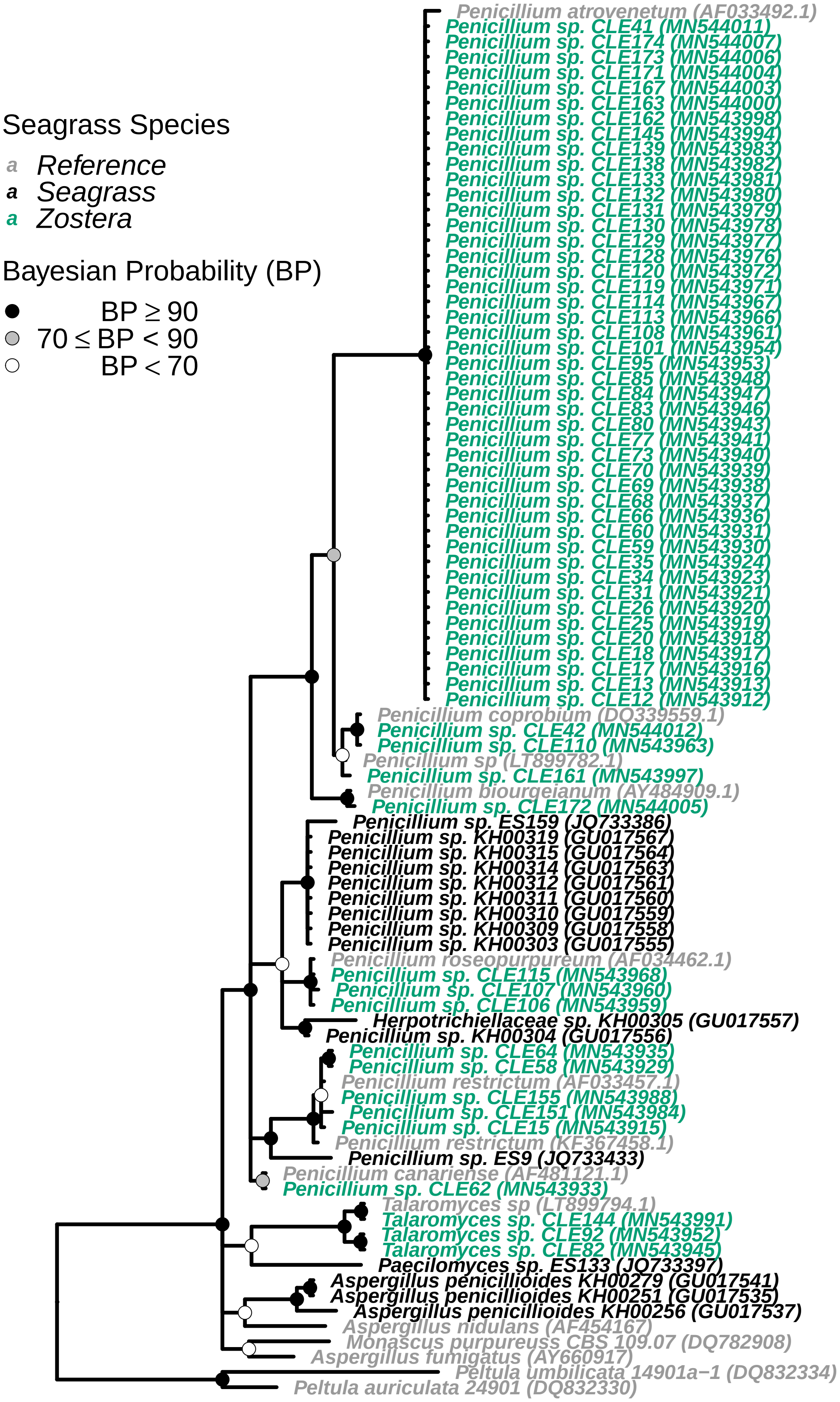
Seagrass Species
a Reference
a Seagrass
a Zostera
Bayesian Probability (BP)
BP ≥ 90
70 ≤ BP < 90
BP < 70
Penicillium atrovenetum (AF033492.1)
Penicillium sp. CLE41 (MN544011)
Penicillium sp. CLE174 (MN544007)
Penicillium sp. CLE173 (MN544006)
Penicillium sp. CLE171 (MN544004)
Penicillium sp. CLE167 (MN544003)
Penicillium sp. CLE163 (MN544000)
Penicillium sp. CLE162 (MN543998)
Penicillium sp. CLE145 (MN543994)
Penicillium sp. CLE139 (MN543983)
Penicillium sp. CLE138 (MN543982)
Penicillium sp. CLE133 (MN543981)
Penicillium sp. CLE132 (MN543980)
Penicillium sp. CLE131 (MN543979)
Penicillium sp. CLE130 (MN543978)
Penicillium sp. CLE129 (MN543977)
Penicillium sp. CLE128 (MN543976)
Penicillium sp. CLE120 (MN543972)
Penicillium sp. CLE119 (MN543971)
Penicillium sp. CLE114 (MN543967)
Penicillium sp. CLE113 (MN543966)
Penicillium sp. CLE108 (MN543961)
Penicillium sp. CLE101 (MN543954)
Penicillium sp. CLE95 (MN543953)
Penicillium sp. CLE85 (MN543948)
Penicillium sp. CLE84 (MN543947)
Penicillium sp. CLE83 (MN543946)
Penicillium sp. CLE80 (MN543943)
Penicillium sp. CLE77 (MN543941)
Penicillium sp. CLE73 (MN543940)
Penicillium sp. CLE70 (MN543939)
Penicillium sp. CLE69 (MN543938)
Penicillium sp. CLE68 (MN543937)
Penicillium sp. CLE66 (MN543936)
Penicillium sp. CLE60 (MN543931)
Penicillium sp. CLE59 (MN543930)
Penicillium sp. CLE35 (MN543924)
Penicillium sp. CLE34 (MN543923)
Penicillium sp. CLE31 (MN543921)
Penicillium sp. CLE26 (MN543920)
Penicillium sp. CLE25 (MN543919)
Penicillium sp. CLE20 (MN543918)
Penicillium sp. CLE18 (MN543917)
Penicillium sp. CLE17 (MN543916)
Penicillium sp. CLE13 (MN543913)
Penicillium sp. CLE12 (MN543912)
Penicillium coprobium (DQ339559.1)
Penicillium sp. CLE42 (MN544012)
Penicillium sp. CLE110 (MN543963)
Penicillium sp (LT899782.1)
Penicillium sp. CLE161 (MN543997)
Penicillium biourgeianum (AY484909.1)
Penicillium sp. CLE172 (MN544005)
Penicillium sp. ES159 (JQ733386)
Penicillium sp. KH00319 (GU017567)
Penicillium sp. KH00315 (GU017564)
Penicillium sp. KH00314 (GU017563)
Penicillium sp. KH00312 (GU017561)
Penicillium sp. KH00311 (GU017560)
Penicillium sp. KH00310 (GU017559)
Penicillium sp. KH00309 (GU017558)
Penicillium sp. KH00303 (GU017555)
Penicillium roseopurpureum (AF034462.1)
Penicillium sp. CLE115 (MN543968)
Penicillium sp. CLE107 (MN543960)
Penicillium sp. CLE106 (MN543959)
Herpotrichiellaceae sp. KH00305 (GU017557)
Penicillium sp. KH00304 (GU017556)
Penicillium sp. CLE64 (MN543935)
Penicillium sp. CLE58 (MN543929)
Penicillium restrictum (AF033457.1)
Penicillium sp. CLE155 (MN543988)
Penicillium sp. CLE151 (MN543984)
Penicillium sp. CLE15 (MN543915)
Penicillium restrictum (KF367458.1)
Penicillium sp. ES9 (JQ733433)
Penicillium canariense (AF481121.1)
Penicillium sp. CLE62 (MN543933)
Talaromyces sp (LT899794.1)
Talaromyces sp. CLE144 (MN543991)
Talaromyces sp. CLE92 (MN543952)
Talaromyces sp. CLE82 (MN543945)
Paecilomyces sp. ES133 (JQ733397)
Aspergillus penicillioides KH00279 (GU017541)
Aspergillus penicillioides KH00251 (GU017535)
Aspergillus penicillioides KH00256 (GU017537)
Aspergillus nidulans (AF454167)
Monascus purpureuss CBS 109.07 (DQ782908)
Aspergillus fumigatus (AY660917)
Peltula umbilicata 14901a−1 (DQ832334)
Peltula auriculata 24901 (DQ832330)

```
# ggsave(filename = 'Euro_ZM.pdf', plot = last_plot(), device
# = 'pdf', width = 15, height = 20, dpi = 300)
```

##### Visualizing the Sordariomycetes tree

```
# Read in the rooted tree file and the csv containing the
# mapping information (e.g. tip labels, seagrass species,
# tissue isolated from, etc)
sord <- read.mrbayes("sord_rooted.tre")
sord_meta <- read.csv("sord_data.csv")

# Join metadata with the tree
sord_v2 <- full_join(sord, sord_meta, by = "label")

# Plot the phylogeny and then save as a pdf
p = ggtree(sord_v2, color = "black", size = 1.5, linetype = 1) +
    geom_tiplab(aes(label = Tree_Name2, color = SeagrassREF2),
        fontface = "bold.italic", size = 6, offset = 0.1)
p = p + theme(legend.position = c(0.8, 0.6), legend.text = element_text(size = 24,
    face = "italic"), legend.title = element_text(size = 24)) +
    guides(color = guide_legend(title = "Seagrass Species"))
p = p + xlim(0, 6) + scale_color_manual(values = c(Zostera = "#009E73",
    Reference = "#999999", Seagrass = "#000000")) + geom_point2(aes(subset = !isTip &
    !is.na(as.numeric(prob_percent)), fill = cut(as.numeric(prob_percent),
    c(0, 70, 90, 100))), shape = 21, size = 5) + scale_fill_manual(values = c("white",
    "grey", "black"), guide = "legend", name = "Bayesian Probability (BP)",
    breaks = c("(90,100]", "(70,90]", "(0,70]"), labels = expression(BP >=
        90, 70 <= BP * " < 90", BP < 70))
p
```

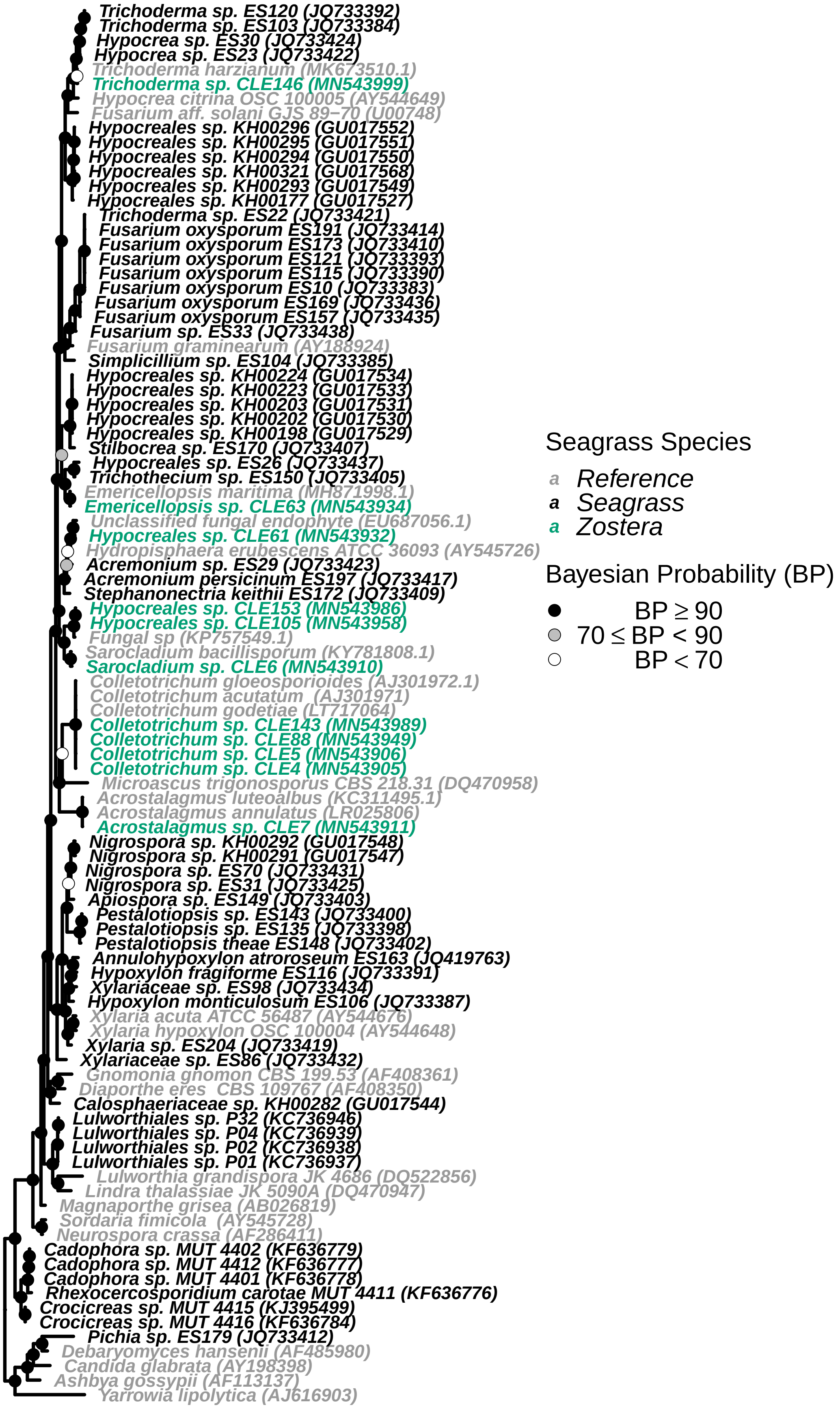

Trichoderma sp. ES120 (JQ733392)
Trichoderma sp. ES103 (JQ733384)
Hypocrea sp. ES30 (JQ733424)
Hypocrea sp. ES23 (JQ733422)
Trichoderma harzianum (MK673510.1)
Trichoderma sp. CLE146 (MN543999)
Hypocrea citrina OSC 100005 (AY544649)
Fusarium aff. solani GJS 89−70 (U00748)
Hypocreales sp. KH00296 (GU017552)
Hypocreales sp. KH00295 (GU017551)
Hypocreales sp. KH00294 (GU017550)
Hypocreales sp. KH00321 (GU017568)
Hypocreales sp. KH00293 (GU017549)
Hypocreales sp. KH00177 (GU017527)
Trichoderma sp. ES22 (JQ733421)
Fusarium oxysporum ES191 (JQ733414)
Fusarium oxysporum ES173 (JQ733410)
Fusarium oxysporum ES121 (JQ733393)
Fusarium oxysporum ES115 (JQ733390)
Fusarium oxysporum ES10 (JQ733383)
Fusarium oxysporum ES169 (JQ733436)
Fusarium oxysporum ES157 (JQ733435)
Fusarium sp. ES33 (JQ733438)
Fusarium graminearum (AY188924)
Simplicillium sp. ES104 (JQ733385)
Hypocreales sp. KH00224 (GU017534)
Hypocreales sp. KH00223 (GU017533)
Hypocreales sp. KH00203 (GU017531)
Hypocreales sp. KH00202 (GU017530)
Hypocreales sp. KH00198 (GU017529)
Stilbocrea sp. ES170 (JQ733407)
Hypocreales sp. ES26 (JQ733437)
Trichothecium sp. ES150 (JQ733405)
Emericellopsis maritima (MH871998.1)
Emericellopsis sp. CLE63 (MN543934)
Unclassified fungal endophyte (EU687056.1)
Hypocreales sp. CLE61 (MN543932)
Hydropisphaera erubescens ATCC 36093 (AY545726)
Acremonium sp. ES29 (JQ733423)
Acremonium persicinum ES197 (JQ733417)
Stephanonectria keithii ES172 (JQ733409)
Hypocreales sp. CLE153 (MN543986)
Hypocreales sp. CLE105 (MN543958)
Fungal sp (KP757549.1)
Sarocladium bacillisporum (KY781808.1)
Sarocladium sp. CLE6 (MN543910)
Colletotrichum gloeosporioides (AJ301972.1)
Colletotrichum acutatum (AJ301971)
Colletotrichum godetiae (LT717064)
Colletotrichum sp. CLE143 (MN543989)
Colletotrichum sp. CLE88 (MN543949)
Colletotrichum sp. CLE5 (MN543906)
Colletotrichum sp. CLE4 (MN543905)
Microascus trigonosporus CBS 218.31 (DQ470958)
Acrostalagmus luteoalbus (KC311495.1)
Acrostalagmus annulatus (LR025806)
Acrostalagmus sp. CLE7 (MN543911)
Nigrospora sp. KH00292 (GU017548)
Nigrospora sp. KH00291 (GU017547)
Nigrospora sp. ES70 (JQ733431)
Nigrospora sp. ES31 (JQ733425)
Apiospora sp. ES149 (JQ733403)
Pestalotiopsis sp. ES143 (JQ733400)
Pestalotiopsis sp. ES135 (JQ733398)
Pestalotiopsis theae ES148 (JQ733402)
Annulohypoxylon atroroseum ES163 (JQ419763)
Hypoxylon fragiforme ES116 (JQ733391)
Xylariaceae sp. ES98 (JQ733434)
Hypoxylon monticulosum ES106 (JQ733387)
Xylaria acuta ATCC 56487 (AY544676)
Xylaria hypoxylon OSC 100004 (AY544648)
Xylaria sp. ES204 (JQ733419)
Xylariaceae sp. ES86 (JQ733432)
Gnomonia gnomon CBS 199.53 (AF408361)
Diaporthe eres CBS 109767 (AF408350)
Calosphaeriaceae sp. KH00282 (GU017544)
Lulworthiales sp. P32 (KC736946)
Lulworthiales sp. P04 (KC736939)
Lulworthiales sp. P02 (KC736938)
Lulworthiales sp. P01 (KC736937)
Lulworthia grandispora JK 4686 (DQ522856)
Lindra thalassiae JK 5090A (DQ470947)
Magnaporthe grisea (AB026819)
Sordaria fimicola (AY545728)
Neurospora crassa (AF286411)
Cadophora sp. MUT 4402 (KF636779)
Cadophora sp. MUT 4412 (KF636777)
Cadophora sp. MUT 4401 (KF636778)
Rhexocercosporidium carotae MUT 4411 (KF636776)
Crocicreas sp. MUT 4415 (KJ395499)
Crocicreas sp. MUT 4416 (KF636784)
Pichia sp. ES179 (JQ733412)
Debaryomyces hansenii (AF485980)
Candida glabrata (AY198398)
Ashbya gossypii (AF113137)
Yarrowia lipolytica (AJ616903)
Seagrass Species
a Reference
a Seagrass
a Zostera
Bayesian Probability (BP)
BP ≥ 90
70 ≤ BP < 90
BP < 70

```
# ggsave(filename = 'Sord_ZM.pdf', plot = last_plot(), device
# = 'pdf', width = 15, height = 25, dpi = 300)
```

##### Visualizing the Dothideomycetes tree

```
# Read in the rooted tree file and the csv containing the
# mapping information (e.g. tip labels, seagrass species,
# tissue isolated from, etc)
doth <- read.mrbayes("doth_rooted.tre")
doth_meta <- read.csv("doth_data.csv")

# Join metadata with the tree
doth_v2 <- full_join(doth, doth_meta, by = "label")

# Plot the phylogeny and then save as a pdf
p = ggtree(doth_v2, color = "black", size = 1.5, linetype = 1) +
    geom_tiplab(aes(label = Tree_Name2, color = SeagrassREF2),
        fontface = "bold.italic", size = 6, offset = 0.1)
p = p + theme(legend.position = c(0.85, 0.575), legend.text = element_text(size = 24,
    face = "italic"), legend.title = element_text(size = 24)) +
    guides(color = guide_legend(title = "Seagrass Species"))
p = p + xlim(0, 6) + scale_color_manual(values = c(Zostera = "#009E73",
    Reference = "#999999", Seagrass = "#000000")) + geom_point2(aes(subset = !isTip &
    !is.na(as.numeric(prob_percent)), fill = cut(as.numeric(prob_percent),
    c(0, 70, 90, 100))), shape = 21, size = 5) + scale_fill_manual(values = c("white",
    "grey", "black"), guide = "legend", name = "Bayesian Probability (BP)",
    breaks = c("(90,100]", "(70,90]", "(0,70]"), labels = expression(BP >=
        90, 70 <= BP * " < 90", BP < 70))
p
```

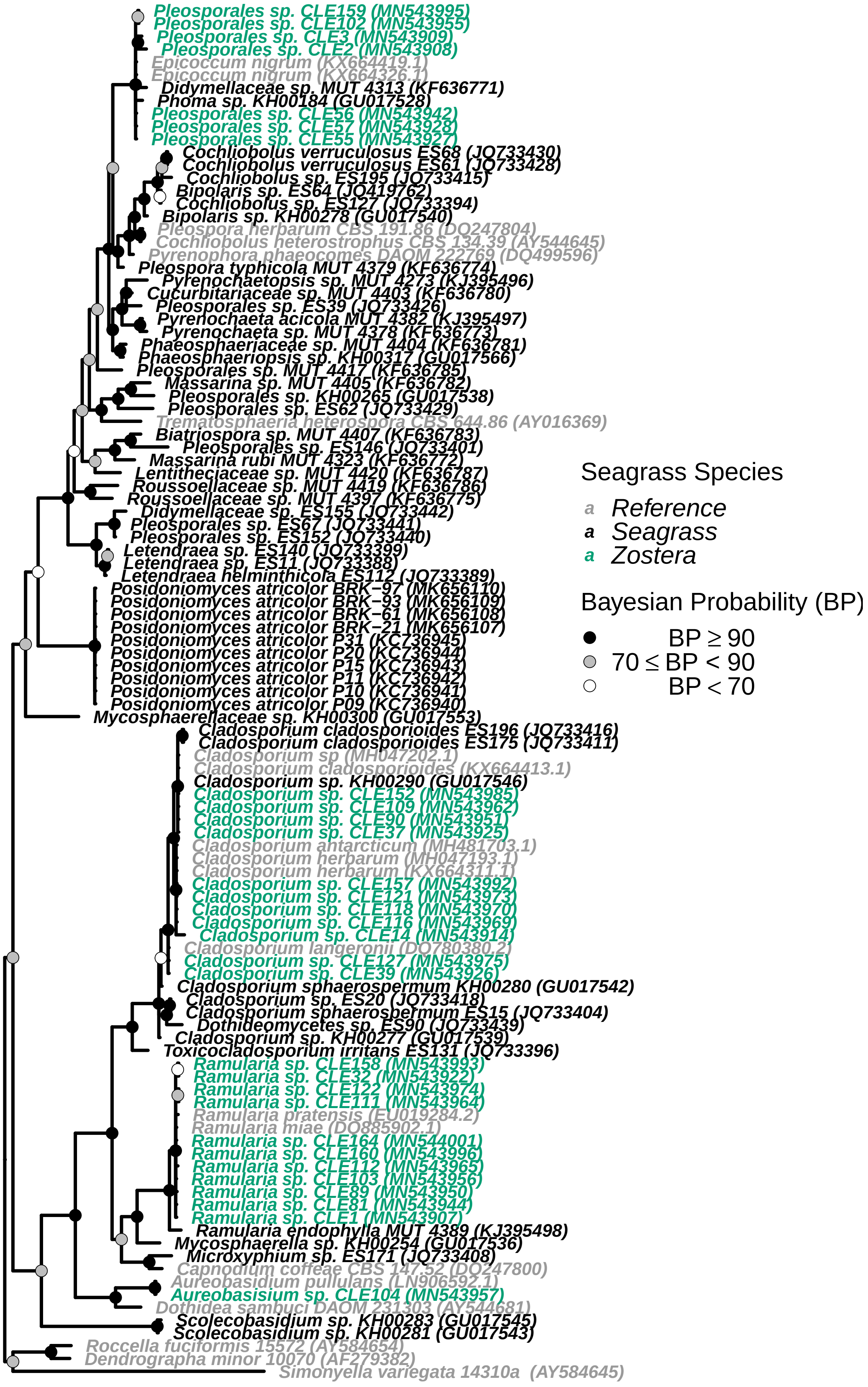
Pleosporales sp. CLE159 (MN543995)
Pleosporales sp. CLE102 (MN543955)
Pleosporales sp. CLE3 (MN543909)
Pleosporales sp. CLE2 (MN543908)
Epicoccum nigrum (KX664419.1)
Epicoccum nigrum (KX664326.1)
Didymellaceae sp. MUT 4313 (KF636771)
Phoma sp. KH00184 (GU017528)
Pleosporales sp. CLE56 (MN543942)
Pleosporales sp. CLE57 (MN543928)
Pleosporales sp. CLE55 (MN543927)
Cochliobolus verruculosus ES68 (JQ733430)
Cochliobolus verruculosus ES61 (JQ733428)
Cochliobolus sp. ES195 (JQ733415)
Bipolaris sp. ES64 (JQ419762)
Cochliobolus sp. ES127 (JQ733394)
Bipolaris sp. KH00278 (GU017540)
Pleospora herbarum CBS 191.86 (DQ247804)
Cochliobolus heterostrophus CBS 134.39 (AY544645)
Pyrenophora phaeocomes DAOM 222769 (DQ499596)
Pleospora typhicola MUT 4379 (KF636774)
Pyrenochaetopsis sp. MUT 4273 (KJ395496)
Cucurbitariaceae sp. MUT 4403 (KF636780)
Pleosporales sp. ES39 (JQ733426)
Pyrenochaeta acicola MUT 4382 (KJ395497)
Pyrenochaeta sp. MUT 4378 (KF636773)
Phaeosphaeriaceae sp. MUT 4404 (KF636781)
Phaeosphaeriopsis sp. KH00317 (GU017566)
Pleosporales sp. MUT 4417 (KF636785)
Massarina sp. MUT 4405 (KF636782)
Pleosporales sp. KH00265 (GU017538)
Pleosporales sp. ES62 (JQ733429)
Trematosphaeria heterospora CBS 644.86 (AY016369)
Biatriospora sp. MUT 4407 (KF636783)
Pleosporales sp. ES146 (JQ733401)
Massarina rubi MUT 4323 (KF636772)
Lentitheciaceae sp. MUT 4420 (KF636787)
Roussoellaceae sp. MUT 4419 (KF636786)
Roussoellaceae sp. MUT 4397 (KF636775)
Didymellaceae sp. ES155 (JQ733442)
Pleosporales sp. ES67 (JQ733441)
Pleosporales sp. ES152 (JQ733440)
Letendraea sp. ES140 (JQ733399)
Letendraea sp. ES11 (JQ733388)
Letendraea helminthicola ES112 (JQ733389)
Posidoniomyces atricolor BRK−97 (MK656110)
Posidoniomyces atricolor BRK−93 (MK656109)
Posidoniomyces atricolor BRK−61 (MK656108)
Posidoniomyces atricolor BRK−21 (MK656107)
Posidoniomyces atricolor P31 (KC736945)
Posidoniomyces atricolor P20 (KC736944)
Posidoniomyces atricolor P15 (KC736943)
Posidoniomyces atricolor P11 (KC736942)
Posidoniomyces atricolor P10 (KC736941)
Posidoniomyces atricolor P09 (KC736940)
Mycosphaerellaceae sp. KH00300 (GU017553)
Cladosporium cladosporioides ES196 (JQ733416)
Cladosporium cladosporioides ES175 (JQ733411)
Cladosporium sp (MH047202.1)
Cladosporium cladosporioides (KX664413.1)
Cladosporium sp. KH00290 (GU017546)
Cladosporium sp. CLE152 (MN543985)
Cladosporium sp. CLE109 (MN543962)
Cladosporium sp. CLE90 (MN543951)
Cladosporium sp. CLE37 (MN543925)
Cladosporium antarcticum (MH481703.1)
Cladosporium herbarum (MH047193.1)
Cladosporium herbarum (KX664311.1)
Cladosporium sp. CLE157 (MN543992)
Cladosporium sp. CLE121 (MN543973)
Cladosporium sp. CLE118 (MN543970)
Cladosporium sp. CLE116 (MN543969)
Cladosporium sp. CLE14 (MN543914)
Cladosporium langeronii (DQ780380.2)
Cladosporium sp. CLE127 (MN543975)
Cladosporium sp. CLE39 (MN543926)
Cladosporium sphaerospermum KH00280 (GU017542)
Cladosporium sp. ES20 (JQ733418)
Cladosporium sphaerospermum ES15 (JQ733404)
Dothideomycetes sp. ES90 (JQ733439)
Cladosporium sp. KH00277 (GU017539)
Toxicocladosporium irritans ES131 (JQ733396)
Ramularia sp. CLE158 (MN543993)
Ramularia sp. CLE32 (MN543922)
Ramularia sp. CLE122 (MN543974)
Ramularia sp. CLE111 (MN543964)
Ramularia pratensis (EU019284.2)
Ramularia miae (DQ885902.1)
Ramularia sp. CLE164 (MN544001)
Ramularia sp. CLE160 (MN543996)
Ramularia sp. CLE112 (MN543965)
Ramularia sp. CLE103 (MN543956)
Ramularia sp. CLE89 (MN543950)
Ramularia sp. CLE81 (MN543944)
Ramularia sp. CLE1 (MN543907)
Ramularia endophylla MUT 4389 (KJ395498)
Mycosphaerella sp. KH00254 (GU017536)
Microxyphium sp. ES171 (JQ733408)
Capnodium coffeae CBS 147.52 (DQ247800)
Aureobasidium pullulans (LN906592.1)
Aureobasisium sp. CLE104 (MN543957)
Dothidea sambuci DAOM 231303 (AY544681)
Scolecobasidium sp. KH00283 (GU017545)
Scolecobasidium sp. KH00281 (GU017543)
Roccella fuciformis 15572 (AY584654)
Dendrographa minor 10070 (AF279382)
Simonyella variegata 14310a (AY584645)
Seagrass Species
a Reference
a Seagrass
a Zostera
Bayesian Probability (BP)
BP ≥ 90
70 ≤ BP < 90
BP < 70

```r
# ggsave(filename = 'Doth_ZM.pdf', plot = last_plot(), device
# = 'pdf', width = 15, height = 25, dpi = 300)
```

#### Visualizing the Basidiomycota and Zygomycota tree

```r
# Read in the rooted tree file and the csv containing the
# mapping information (e.g. tip labels, seagrass species,
# tissue isolated from, etc)
bz <- read.mrbayes("bz_root.tre")
bz_meta <- read.csv("bz_data.csv")

# Join metadata with the tree
bz_v2 <- full_join(bz, bz_meta, by = "label")

# Plot the phylogeny and then save as a pdf
p = ggtree(bz_v2, color = "black", size = 1.5, linetype = 1) +
    geom_tiplab(aes(label = Tree_Name2, color = SeagrassREF2),
        fontface = "bold.italic", size = 6, offset = 0.1)
p = p + theme(legend.position = c(0.8, 0.6), legend.text = element_text(size = 24,
    face = "italic"), legend.title = element_text(size = 24)) +
    guides(color = guide_legend(title = "Seagrass Species"))
p = p + xlim(0, 6) + scale_color_manual(values = c(Zostera = "#009E73",
    Reference = "#999999", Seagrass = "#000000")) + geom_point2(aes(subset = !isTip &
    !is.na(as.numeric(prob_percent)), fill = cut(as.numeric(prob_percent),
    c(0, 70, 90, 100))), shape = 21, size = 5) + scale_fill_manual(values = c("white",
    "grey", "black"), guide = "legend", name = "Bayesian Probability (BP)",
    breaks = c("(90,100]", "(70,90]", "(0,70]"), labels = expression(BP >=
        90, 70 <= BP * " < 90", BP < 70))
p
```

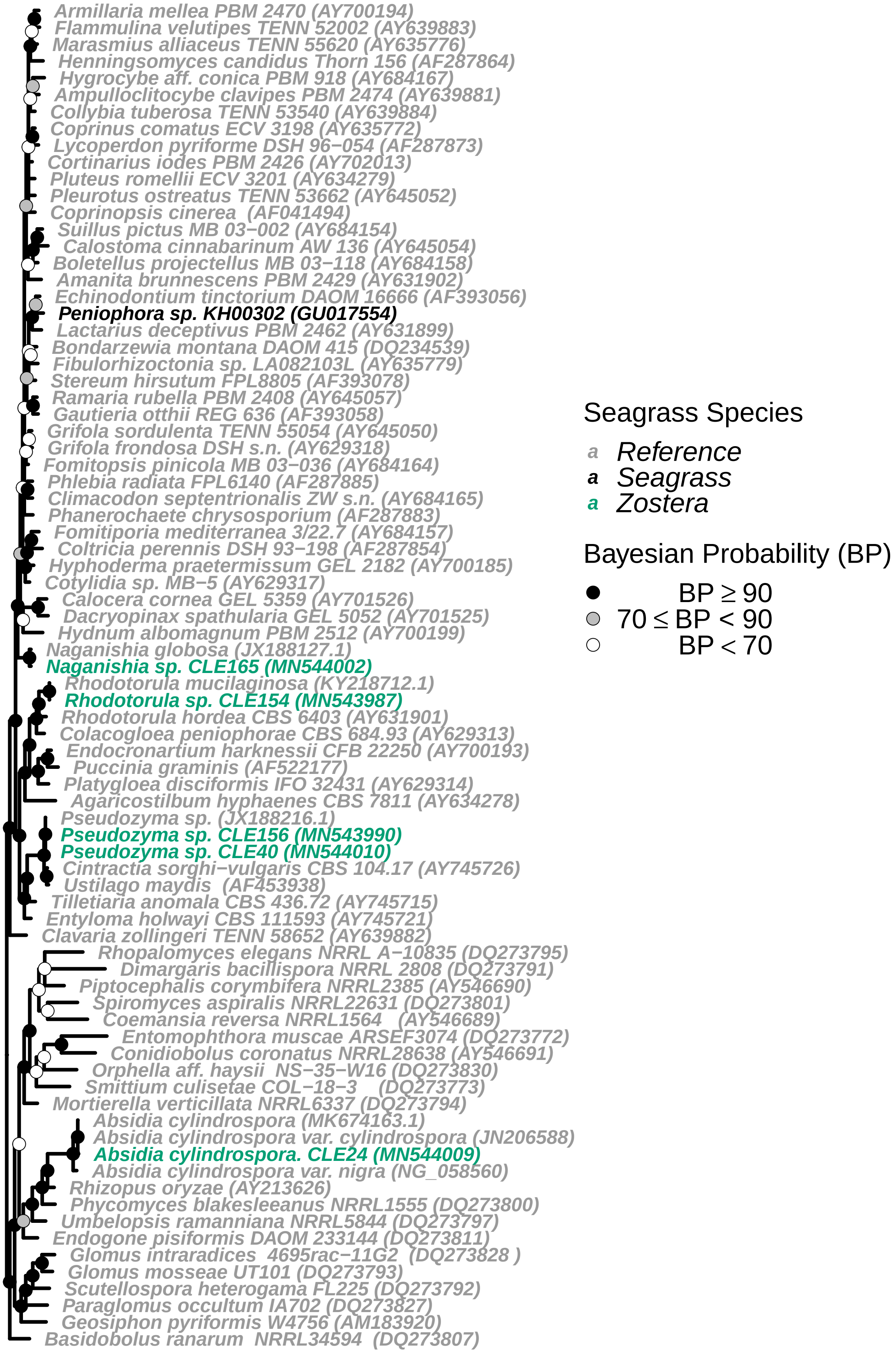
Armillaria mellea PBM 2470 (AY700194)
Flammulina velutipes TENN 52002 (AY639883)
Marasmius alliaceus TENN 55620 (AY635776)
Henningsomyces candidus Thorn 156 (AF287864)
Hygrocybe aff. conica PBM 918 (AY684167)
Ampulloclitocybe clavipes PBM 2474 (AY639881)
Collybia tuberosa TENN 53540 (AY639884)
Coprinus comatus ECV 3198 (AY635772)
Lycoperdon pyriforme DSH 96−054 (AF287873)
Cortinarius iodes PBM 2426 (AY702013)
Pluteus romellii ECV 3201 (AY634279)
Pleurotus ostreatus TENN 53662 (AY645052)
Coprinopsis cinerea (AF041494)
Suillus pictus MB 03−002 (AY684154)
Calostoma cinnabarinum AW 136 (AY645054)
Boletellus projectellus MB 03−118 (AY684158)
Amanita brunnescens PBM 2429 (AY631902)
Echinodontium tinctorium DAOM 16666 (AF393056)
Peniophora sp. KH00302 (GU017554)
Lactarius deceptivus PBM 2462 (AY631899)
Bondarzewia montana DAOM 415 (DQ234539)
Fibulorhizoctonia sp. LA082103L (AY635779)
Stereum hirsutum FPL8805 (AF393078)
Ramaria rubella PBM 2408 (AY645057)
Gautieria otthii REG 636 (AF393058)
Grifola sordulenta TENN 55054 (AY645050)
Grifola frondosa DSH s.n. (AY629318)
Fomitopsis pinicola MB 03−036 (AY684164)
Phlebia radiata FPL6140 (AF287885)
Climacodon septentrionalis ZW s.n. (AY684165)
Phanerochaete chrysosporium (AF287883)
Fomitiporia mediterranea 3/22.7 (AY684157)
Coltricia perennis DSH 93−198 (AF287854)
Hyphoderma praetermissum GEL 2182 (AY700185)
Cotylidia sp. MB−5 (AY629317)
Calocera cornea GEL 5359 (AY701526)
Dacryopinax spathularia GEL 5052 (AY701525)
Hydnum albomagnum PBM 2512 (AY700199)
Naganishia globosa (JX188127.1)
Naganishia sp. CLE165 (MN544002)
Rhodotorula mucilaginosa (KY218712.1)
Rhodotorula sp. CLE154 (MN543987)
Rhodotorula hordea CBS 6403 (AY631901)
Colacogloea peniophorae CBS 684.93 (AY629313)
Endocronartium harknessii CFB 22250 (AY700193)
Puccinia graminis (AF522177)
Platygloea disciformis IFO 32431 (AY629314)
Agaricostilbum hyphaenes CBS 7811 (AY634278)
Pseudozyma sp. (JX188216.1)
Pseudozyma sp. CLE156 (MN543990)
Pseudozyma sp. CLE40 (MN544010)
Cintractia sorghi−vulgaris CBS 104.17 (AY745726)
Ustilago maydis (AF453938)
Tilletiaria anomala CBS 436.72 (AY745715)
Entyloma holwayi CBS 111593 (AY745721)
Clavaria zollingeri TENN 58652 (AY639882)
Rhopalomyces elegans NRRL A−10835 (DQ273795)
Dimargaris bacillispora NRRL 2808 (DQ273791)
Piptocephalis corymbifera NRRL2385 (AY546690)
Spiromyces aspiralis NRRL22631 (DQ273801)
Coemansia reversa NRRL1564 (AY546689)
Entomophthora muscae ARSEF3074 (DQ273772)
Conidiobolus coronatus NRRL28638 (AY546691)
Orphella aff. haysii NS−35−W16 (DQ273830)
Smittium culisetae COL−18−3 (DQ273773)
Mortierella verticillata NRRL6337 (DQ273794)
Absidia cylindrospora (MK674163.1)
Absidia cylindrospora var. cylindrospora (JN206588)
Absidia cylindrospora. CLE24 (MN544009)
Absidia cylindrospora var. nigra (NG_058560)
Rhizopus oryzae (AY213626)
Phycomyces blakesleeanus NRRL1555 (DQ273800)
Umbelopsis ramanniana NRRL5844 (DQ273797)
Endogone pisiformis DAOM 233144 (DQ273811)
Glomus intraradices 4695rac−11G2 (DQ273828 )
Glomus mosseae UT101 (DQ273793)
Scutellospora heterogama FL225 (DQ273792)
Paraglomus occultum IA702 (DQ273827)
Geosiphon pyriformis W4756 (AM183920)
Basidobolus ranarum NRRL34594 (DQ273807)
Seagrass Species
a Reference
a Seagrass
a Zostera
Bayesian Probability (BP)
BP ≥ 90
70 ≤ BP < 90
BP < 70

```
# ggsave(filename = 'Bz_ZM.pdf', plot = last_plot(), device =
# 'pdf', width = 15, height = 20, dpi = 300)
```

##### Distribution of fungal counts across isolation sources

```
# Read in csv containing isolate information
fungi_meta <- read.csv("fungi_tab1.csv")

# Reorder fungal orders to be alphabetical by phylum
fungi_meta$Order_f = factor(fungi_meta$Order, levels = c("Capnodiales",
    "Dothideales", "Pleosporales", "Eurotiales", "Glomerellales",
    "Hypocreales", "Filobasidiales", "Sporidiobolales", "Ustilaginales",
    "Mucorales"))

# Count the number of isolation sources (tissue types) for
# each taxonomic order
fungi_meta2 <- fungi_meta %>% count(Order_f, Tissue)

# Rearange the count values and determine a value to use to
# place the count on each bar in the histogram
fungi_meta3 <- fungi_meta2 %>% group_by(Order_f) %>% arrange(Order_f,
    desc(Tissue)) %>% mutate(lab_ypos = cumsum(n) - 0.5 * n)

# Generate the histogram of fungal counts x isolation sources
p <- ggplot(data = fungi_meta3, aes(x = Order_f, y = n, fill = Tissue)) +
    geom_col() + geom_text(aes(y = lab_ypos, label = n, group = Tissue),
    fontface = "bold", size = 6, color = "white")
p + theme(axis.text.x = element_text(angle = -70, hjust = 0,
    vjust = 0.5)) + theme(text = element_text(size = 34)) + ylab("Number of Isolates") +
    guides(fill = guide_legend(title = "Isolated From")) + scale_fill_viridis_d(option = "C",
    begin = 0.85, end = 0) + xlab("Order")
```

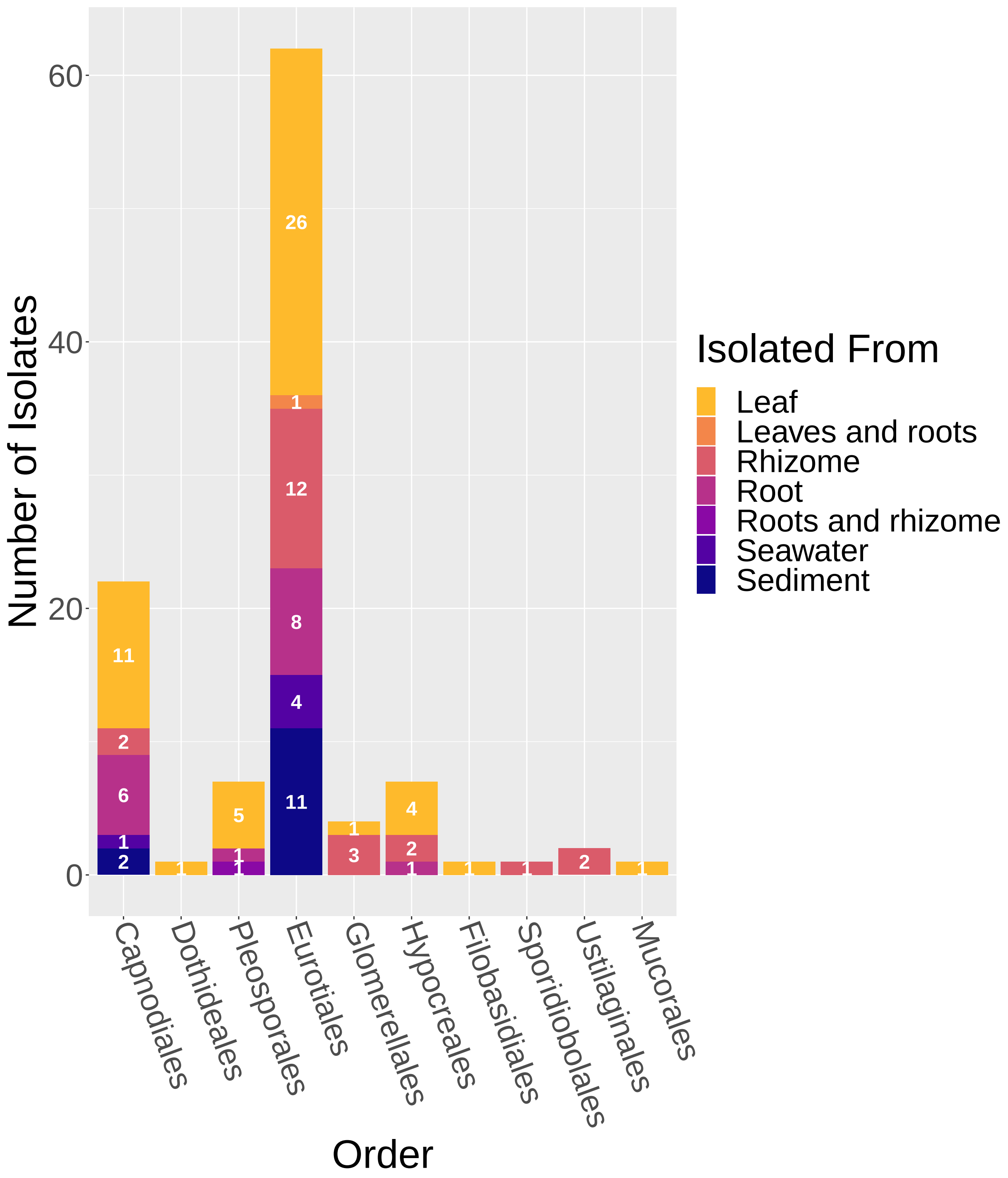


```
# ggsave(filename = 'Fungi_Hist.pdf', plot = last_plot(),
# device = 'pdf', width = 14, height = 18, dpi = 300)
```

##### Distribution of fungal counts across media types

```
# Count the number of media types for each taxonomic order
fungi_meta_media <- fungi_meta %>% count(Order_f, Media_recipe)

# Rearange the count values and determine a value to use to
# place the count on each bar in the histogram
fungi_meta_media <- fungi_meta_media %>% group_by(Order_f) %>%
    arrange(Order_f, desc(Media_recipe)) %>% mutate(lab_ypos = cumsum(n) -
    0.5 * n)

# Generate the histogram of fungal counts x media types
p <- ggplot(data = fungi_meta_media, aes(x = Order_f, y = n,
    fill = Media_recipe)) + geom_col() + geom_text(aes(y = lab_ypos,
    label = n, group = Media_recipe), fontface = "bold", size = 6,
    color = "white")
p + theme(axis.text.x = element_text(angle = -70, hjust = 0,
    vjust = 0.5)) + theme(text = element_text(size = 34)) + ylab("Number of Isolates") +
    guides(fill = guide_legend(title = "Media recipe")) + scale_fill_viridis_d(option = "C",
    begin = 0.85, end = 0) + xlab("Order")
```

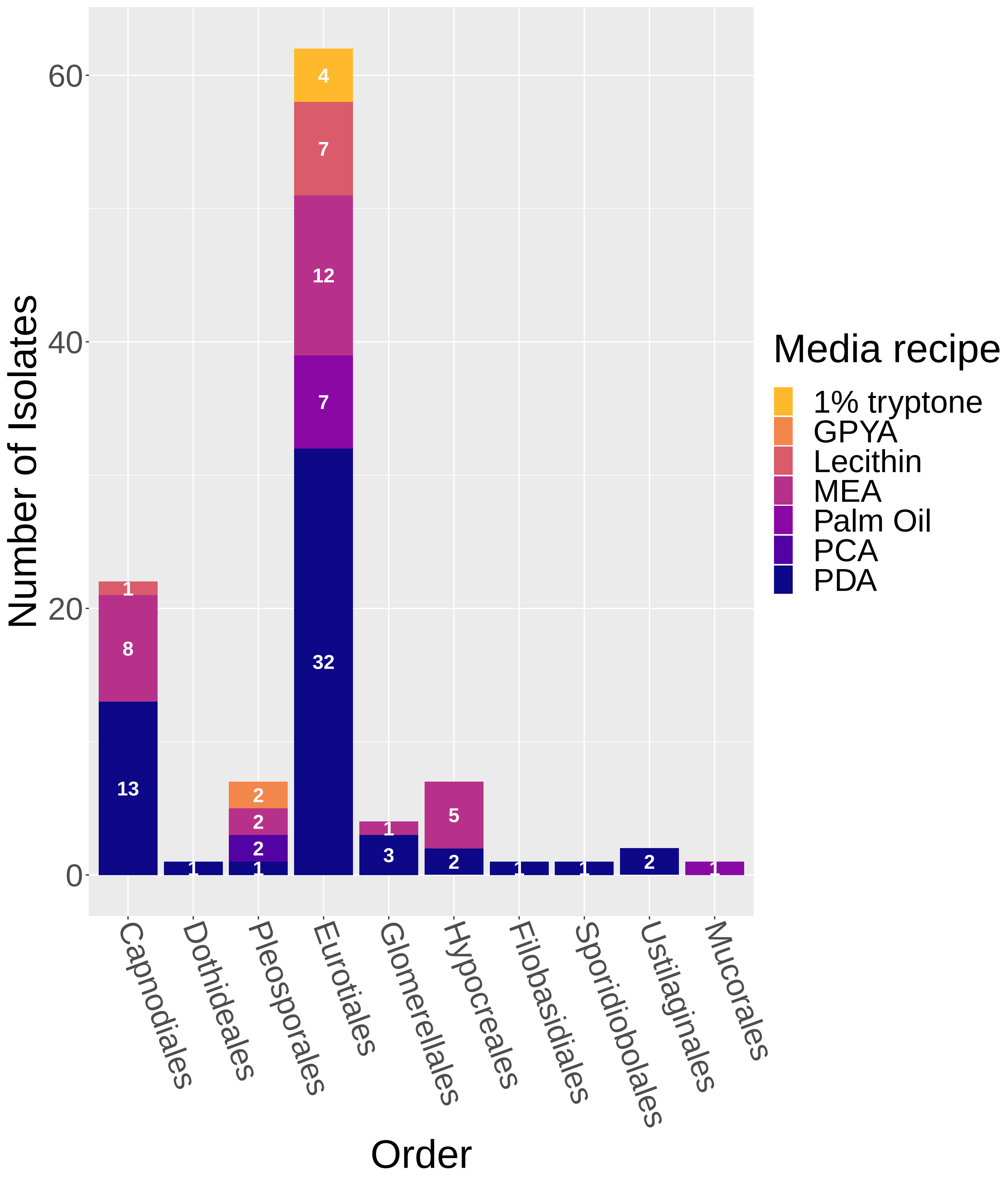


```
# ggsave(filename = 'Fun_Hist_Media.pdf', plot = last_plot(),
# device = 'pdf', width = 14, height = 18, dpi = 300)
```

#### Exploring relationships between fungal genera and where / what they were isolated from / on

```
# Copy fungi_meta for subsetting
fungi_meta_trend <- fungi_meta

# remove media types not frequently used / or not used with
# most tissue types (1% tryptone, PCA, GPYA) remove isolation
# sources not frequently used / or not used with most tissue
# types (seawater, leafs and roots, roots and rhizome)
fungi_meta_trend <- filter(fungi_meta_trend, Tissue != "Seawater")
fungi_meta_trend <- filter(fungi_meta_trend, Tissue != "Leaves and roots")
fungi_meta_trend <- filter(fungi_meta_trend, Tissue != "Roots and rhizome")
fungi_meta_trend <- filter(fungi_meta_trend, Media_recipe !=
    "PCA")
fungi_meta_trend <- filter(fungi_meta_trend, Media_recipe !=
    "1% tryptone")
fungi_meta_trend <- filter(fungi_meta_trend, Media_recipe !=
    "GPYA")
```

#### Is there a relationship between the number of media types a fungal genus was cultured from and the number of isolation sources (tissue types) it was cultured from?

```
# Select appropriate columns
fungi_meta_trend2 <- fungi_meta_trend %>% select(Molecular.ID,
    Tissue, Media_recipe)

# Count the number of different isolation sources (Tissue)
# and media types (Media_recipe) that resulted in the
# isolation of a particular fungal genus
fungi_meta_trend3 <- fungi_meta_trend2 %>% group_by(Molecular.ID) %>%
    summarise(n_tissue = n_distinct(Tissue), n_med = n_distinct(Media_recipe))

# Generate plot
p <- ggplot(data = fungi_meta_trend3, aes(x = n_tissue, y = n_med)) +
    geom_point(size = 5, position = position_jitter(width = 0.1,
        height = 0.05)) + geom_smooth(method = lm)
p <- p + theme(text = element_text(size = 18)) + ylab("Number of media types") +
    xlab("Number of isolation sources")
p
```

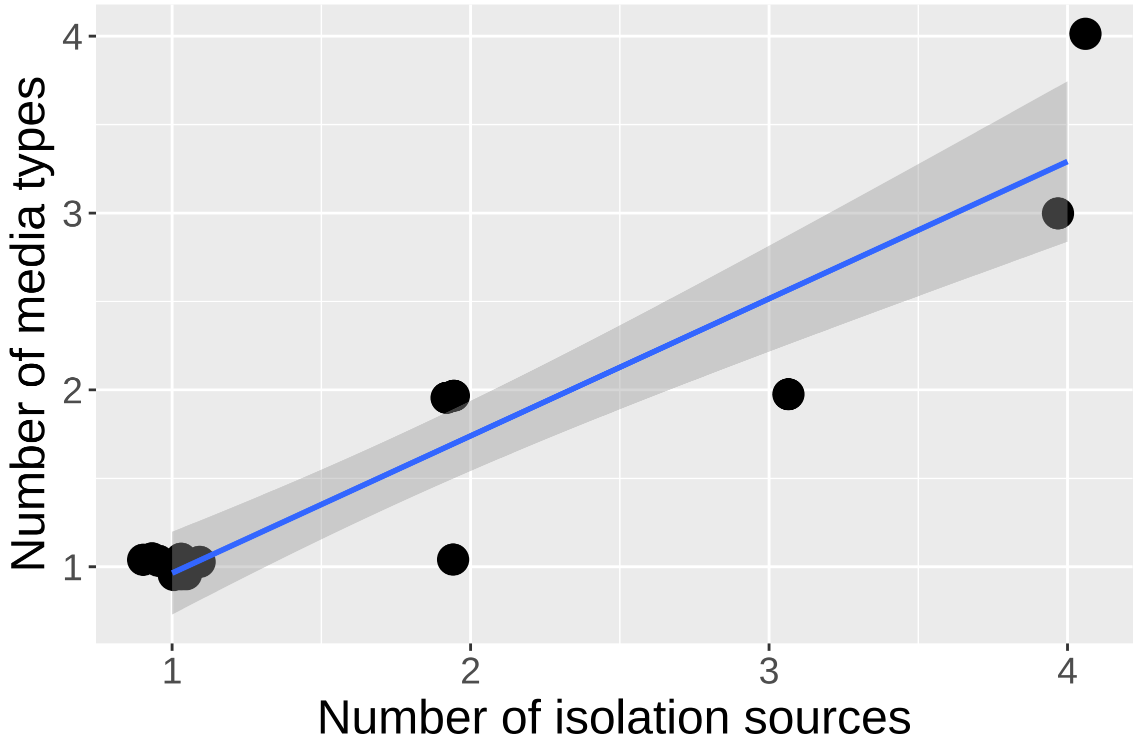


```r
# Linear model
fit1 <- lm(n_tissue ~ n_med, data = fungi_meta_trend3)
summary(fit1)
```

```
##
## Call:
## lm(formula = n_tissue ~ n_med, data = fungi_meta_trend3)
##
## Residuals:
##     Min      1Q  Median      3Q     Max
## -0.4943 -0.1364 -0.1364 -0.1364  0.8636
##
## Coefficients:
##             Estimate Std. Error t value Pr(>|t|)
## (Intercept)  0.01705    0.21417   0.080    0.938
## n_med        1.11932    0.12099   9.251  4.4e-07 ***
## ---
## Signif. codes:  0 '***' 0.001 '**' 0.01 '*' 0.05 '.' 0.1 ' ' 1
##
## Residual standard error: 0.4144 on 13 degrees of freedom
## Multiple R-squared:  0.8681, Adjusted R-squared:  0.858
## F-statistic: 85.58 on 1 and 13 DF,  p-value: 4.405e-07
```

##### Is there a relationship between the number of salt sources (instant ocean at different concentrations, no salt, seawater) a fungal genus was cultured from and the number of isolation sources (tissue types) it was cultured from?

```
# Select appropriate columns
fungi_meta_trend2 <- fungi_meta_trend %>% select(Molecular.ID,
    Tissue, Salt_source)

# Count the number of different isolation sources (Tissue)
# and salt sources (Salt_source) that resulted in the
# isolation of a particular fungal genus
fungi_meta_trend3 <- fungi_meta_trend2 %>% group_by(Molecular.ID) %>%
    summarise(n_tissue = n_distinct(Tissue), n_salt = n_distinct(Salt_source))

# Generate plot
q <- ggplot(data = fungi_meta_trend3, aes(x = n_tissue, y = n_salt)) +
    geom_point(size = 5, position = position_jitter(width = 0.1,
        height = 0.05)) + geom_smooth(method = lm)
q <- q + theme(text = element_text(size = 18)) + ylab("Number of salt sources") +
    xlab("Number of isolation sources")
q
```

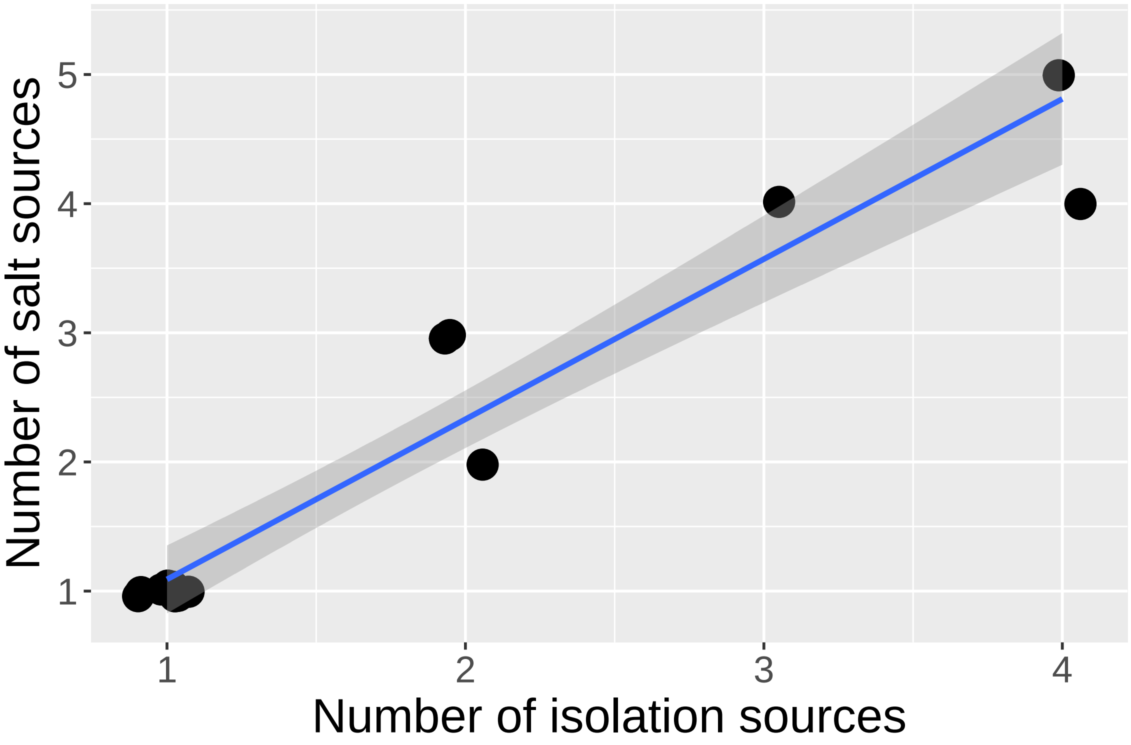


```
# Linear model
fit1 <- lm(n_tissue ~ n_salt, data = fungi_meta_trend3)
summary(fit1)
```

```
##
## Call:
```

```
## lm(formula = n_tissue ~ n_salt, data = fungi_meta_trend3)
##
## Residuals:
##      Min       1Q   Median       3Q      Max
## -0.48333  0.01667  0.01667  0.01667  0.76667
##
## Coefficients:
##             Estimate Std. Error t value Pr(>|t|)
## (Intercept)  0.23333    0.13810    1.69    0.115
## n_salt       0.75000    0.05702   13.15  6.9e-09 ***
## ---
## Signif. codes:  0 '***' 0.001 '**' 0.01 '*' 0.05 '.' 0.1 ' ' 1
##
## Residual standard error: 0.3017 on 13 degrees of freedom
## Multiple R-squared:  0.9301, Adjusted R-squared:  0.9247
## F-statistic:   173 on 1 and 13 DF,  p-value: 6.896e-09
```

#### Is there a relationship between the number of salt sources (instant ocean at different concentrations, no salt, seawater) a fungal genus was cultured from and the number of media recipes it was cultured from?

```r
# Select appropriate columns
fungi_meta_trend2 <- fungi_meta_trend %>% select(Molecular.ID,
    Media_recipe, Salt_source)

# Count the number of different media recipes (Media_recipe)
# and salt sources (Salt_source) that resulted in the
# isolation of a particular fungal genus
fungi_meta_trend3 <- fungi_meta_trend2 %>% group_by(Molecular.ID) %>%
    summarise(n_med = n_distinct(Media_recipe), n_salt = n_distinct(Salt_source))

# Generate plot
r <- ggplot(data = fungi_meta_trend3, aes(x = n_med, y = n_salt)) +
    geom_point(size = 5, position = position_jitter(width = 0.1,
        height = 0.05)) + geom_smooth(method = lm)
r <- r + theme(text = element_text(size = 18)) + ylab("Number of salt sources") +
    xlab("Number of media types")
r
```

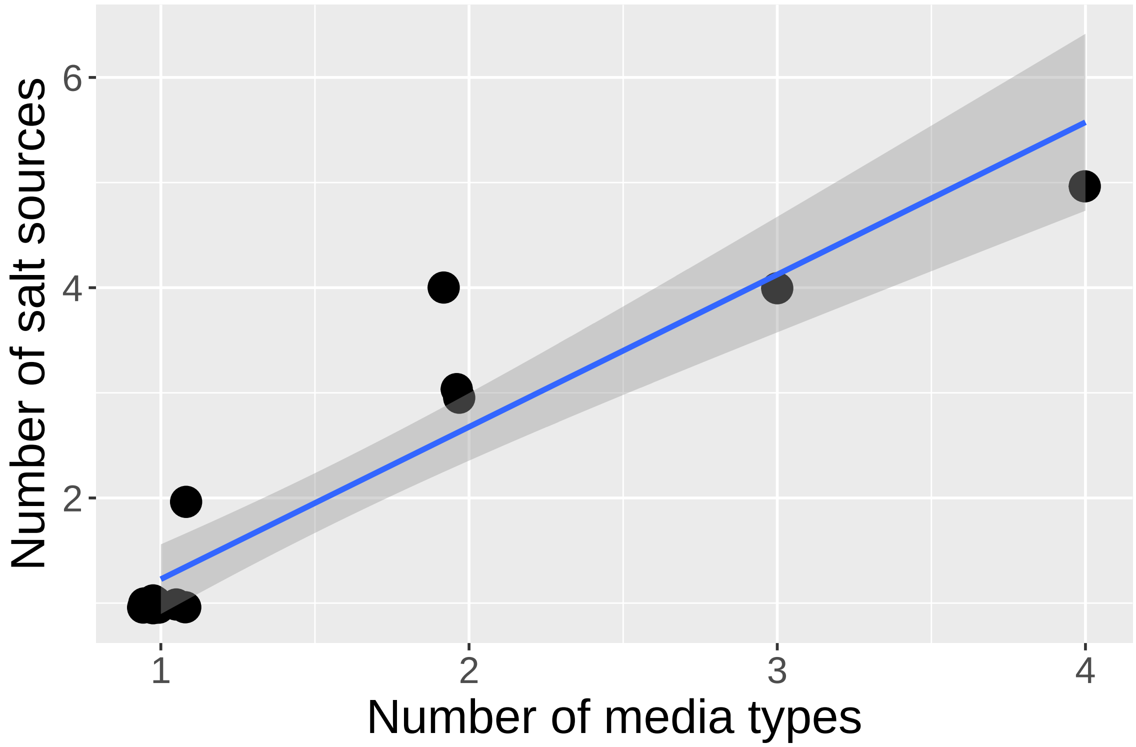


```
# Linear Model
fit1 <- lm(n_med ~ n_salt, data = fungi_meta_trend3)
summary(fit1)
```

```
##
## Call:
## lm(formula = n_med ~ n_salt, data = fungi_meta_trend3)
##
## Residuals:
##      Min       1Q   Median       3Q      Max
## -0.74762 -0.03333  0.07381  0.07381  0.64524
##
## Coefficients:
##             Estimate Std. Error t value Pr(>|t|)
## (Intercept)  0.31905    0.15085   2.115   0.0543 .
## n_salt       0.60714    0.06228   9.749 2.42e-07 ***
## ---
## Signif. codes:  0 '***' 0.001 '**' 0.01 '*' 0.05 '.' 0.1 ' ' 1
##
## Residual standard error: 0.3296 on 13 degrees of freedom
## Multiple R-squared:  0.8797, Adjusted R-squared:  0.8704
## F-statistic: 95.03 on 1 and 13 DF,  p-value: 2.416e-07
```

##### Combine plots for supplemental figure

```
patched <- p + q + r

patched + plot_annotation(tag_levels = "A")
```

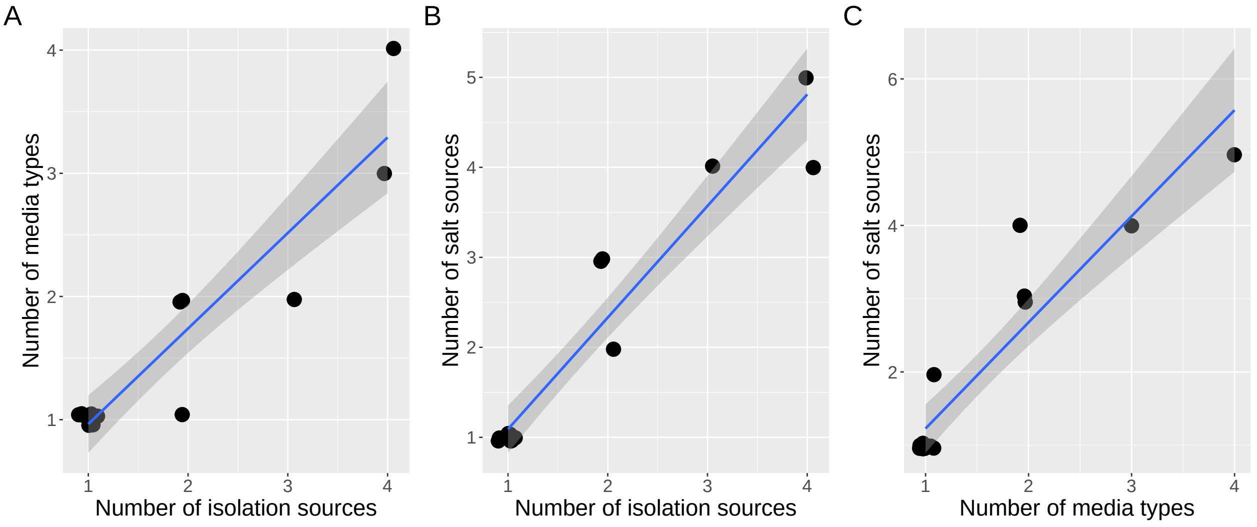


```
# ggsave(filename = 'Fungal_Trends.pdf', plot = last_plot(),
# device = 'pdf', width = 14, height = 4, dpi = 300)
```

##### Distribution of bacterial counts across isolation sources

```
# Read in csv containing isolate information
bact_meta <- read.csv("bact_tab2.csv")

# Reorder bacterial orders to be alphabetical by phylum
bact_meta$Order_f = factor(bact_meta$Order, levels = c("Actinomycetales",
    "Streptomycetales", "Rhizobiales", "Lactobacillales", "Flavobacteriales",
    "Alteromonadales", "Enterobacteriales", "Oceanospirillales",
    "Pseudomonadales", "Vibrionales"))

# Count the number of isolation sources (tissue types) for
# each taxonomic order
bact_meta2 <- bact_meta %>% count(Order_f, Tissue)

# Rearange the count values and determine a value to use to
# place the count on each bar in the histogram
bact_meta3 <- bact_meta2 %>% group_by(Order_f) %>% arrange(Order_f,
    desc(Tissue)) %>% mutate(lab_ypos = cumsum(n) - 0.5 * n)

# Generate the histogram of bacterial counts x isolation
# sources
bac_iso_hist <- ggplot(data = bact_meta3, aes(x = Order_f, y = n,
    fill = Tissue)) + geom_col() + geom_text(aes(y = lab_ypos,
    label = n, group = Tissue), fontface = "bold", size = 7,
    color = "white")
bac_iso_hist <- bac_iso_hist + theme(axis.text.x = element_text(angle = -70,
    hjust = 0, vjust = 0.5)) + theme(text = element_text(size = 28)) +
    ylab("Number of Isolates") + guides(fill = guide_legend(title = "Isolated From")) +
    scale_fill_viridis_d(option = "C", begin = 0.85, end = 0) +
    xlab("Order")
bac_iso_hist
```

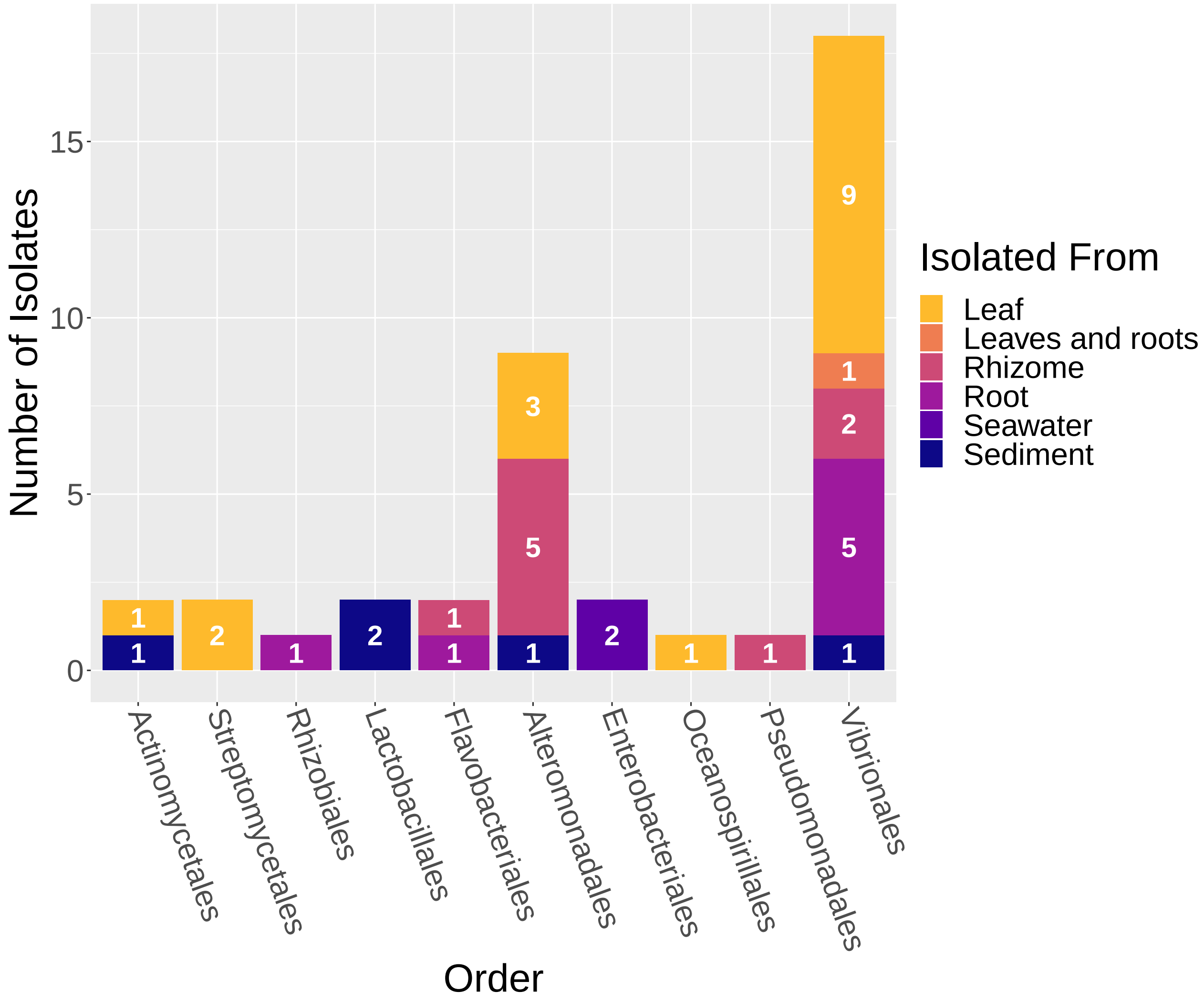


```
# ggsave(filename = 'Bac_Hist.pdf', plot = last_plot(),
# device = 'pdf', width = 10, height = 10, dpi = 300)
```

##### Distribution of bacterial counts across media recipes

```
# Count the number of isolation sources (tissue types) for
# each taxonomic order
bact_meta_media <- bact_meta %>% count(Order_f, Media.recipe)

# Rearange the count values and determine a value to use to
# place the count on each bar in the histogram
bact_meta_media <- bact_meta_media %>% group_by(Order_f) %>%
    arrange(Order_f, desc(Media.recipe)) %>% mutate(lab_ypos = cumsum(n) -
    0.5 * n)

# Generate the histogram of bacterial counts x isolation
# sources
bac_med_hist <- ggplot(data = bact_meta_media, aes(x = Order_f,
    y = n, fill = Media.recipe)) + geom_col() + geom_text(aes(y = lab_ypos,
```

```r
    label = n, group = Media.recipe), fontface = "bold", size = 7,
    color = "white")
bac_med_hist <- bac_med_hist + theme(axis.text.x = element_text(angle = -70,
    hjust = 0, vjust = 0.5)) + theme(text = element_text(size = 28)) +
    ylab("Number of Isolates") + guides(fill = guide_legend(title = "Media recipe")) +
    scale_fill_viridis_d(option = "C", begin = 0.85, end = 0) +
    xlab("Order")
bac_med_hist
```

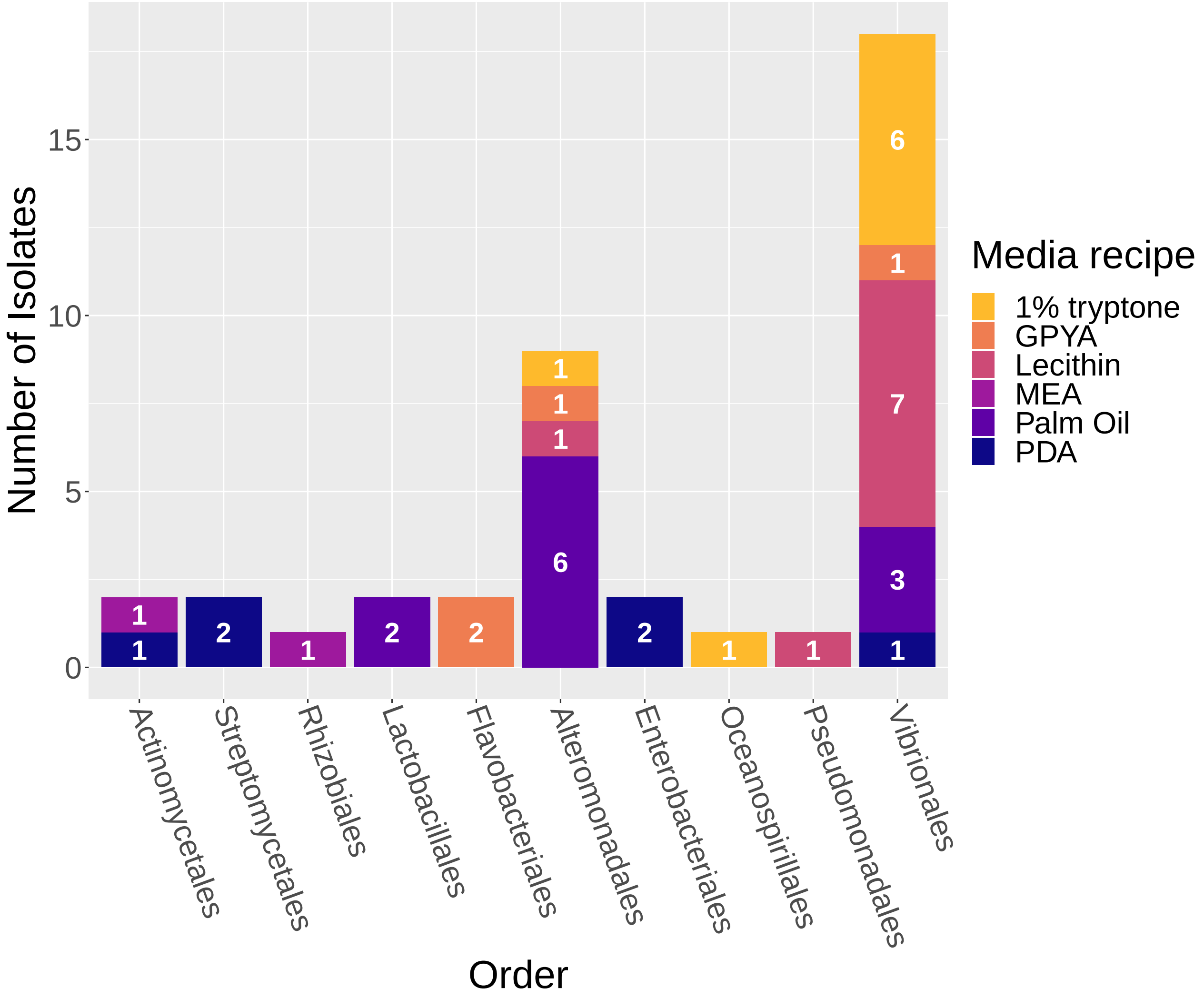


```r
# ggsave(filename = 'Bact_Hist_Media.pdf', plot =
# last_plot(), device = 'pdf', width = 10, height = 10, dpi =
# 300)
```

#### Now combine the two histograms for publication

```r
bac_iso_hist + bac_med_hist + plot_annotation(tag_levels = "A")
```

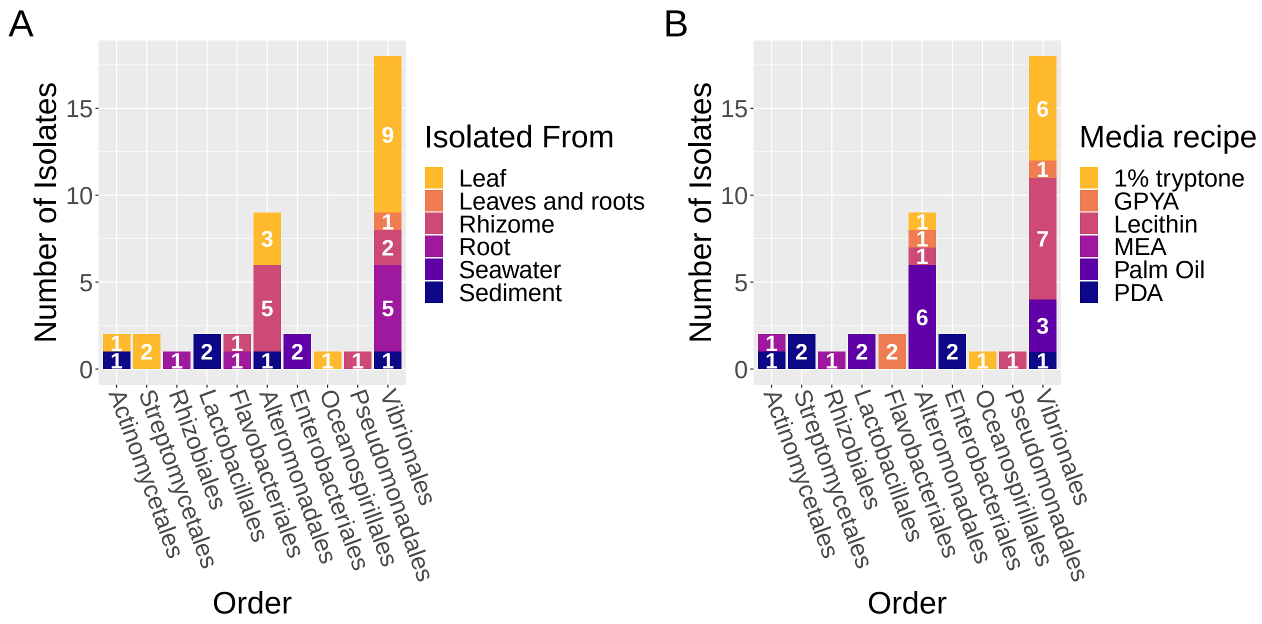


```
# ggsave(filename = 'Bact_Hist_Combined.pdf', plot =
# last_plot(), device = 'pdf', width = 16, height = 8, dpi =
# 300)
```

#### Comparisons to ITS data from Ettinger & Eisen (2019)

##### Obtaining a list of unique genera found in Ettinger & Eisen (2019) to compare to the fungal genera isolated in this study

```
# Import the rarefied subsampled ASV table used in Ettinger &
# Eisen (2019)
SGFungi_rare10000_RA <- readRDS("ps.nocontrols.pear_nomin_err3_rare10000.LOC.NoNEG.RA.noendo.noGP.RDS")

# Convert the phyloseq object to an R dataframe
df <- psmelt(SGFungi_rare10000_RA)

# Get a list of unique genera represented in this ASV table
unique_genera <- unique(df$Genus)

# Note that in the UNITE databse - Mycosphaerella ==
# Ramularia

# Save this list to a csv file write.csv(genus,
# 'SGFungi_rare10000_taxglom_OrderLevel.csv')
```

##### At what relative abudance are the fungal taxa isolated in this study also detected from the same sample types / isolation sources?

```
# combine by order
SGFungi_rare10000_taxglom_OrderLevel_RA <- tax_glom(SGFungi_rare10000_RA,
```

```
    taxrank = "Order", NArm = FALSE)

# Subset phyloseq object to only have the orders isolated
# here (if present)
IsolateOrdersOnly <- subset_taxa(SGFungi_rare10000_taxglom_OrderLevel_RA,
    Order == "o__Capnodiales" | Order == "o__Dothideales" | Order ==
        "o__Pleosporales" | Order == "o__Eurotiales" | Order ==
        "o__Sporidiobolales" | Order == "o__Glomerellales" |
        Order == "o__Hypocreales" | Order == "o__Filobasidiales" |
        Order == "o__Mucorales" | Order == "o__Ustilaginales")

# Convert the phyloseq object to an R dataframe
IsolateOrdersOnly_df <- psmelt(IsolateOrdersOnly)

# Summarize the df and calculate the mean
grouped_order <- group_by(IsolateOrdersOnly_df, SampleType, Order,
    Class, Phylum)
avgs_order <- summarise(grouped_order, mean = 100 * mean(Abundance))

# Order the Orders!
avgs_order$Order_f = factor(avgs_order$Order, levels = c("o__Capnodiales",
    "o__Dothideales", "o__Pleosporales", "o__Eurotiales", "o__Glomerellales",
    "o__Hypocreales", "o__Filobasidiales", "o__Sporidiobolales",
    "o__Ustilaginales", "o__Mucorales"))

# Order the sample types to match the histogram
avgs_order$SampleType <- factor(avgs_order$SampleType, levels = c("Leaf",
    "Rhizome", "Root", "Sediment"))

# Rearange the mean relative abudnance (RA) values and
# determine a value to use to place the RA on each bar in the
# histogram
fungi_order_comp <- avgs_order %>% group_by(Order_f) %>% arrange(Order_f,
    desc(SampleType)) %>% mutate(lab_ypos = cumsum(mean) - 0.5 *
    mean)

fungi_order_comp_2 <- fungi_order_comp %>% mutate(mean_label = round((mean),
    digits = 1))

# Generate the histogram of fungal mean RA x sample type (~
# isolation source) and put the % on the bar if > 1% mean RA
p <- ggplot(data = fungi_order_comp_2, aes(x = Order_f, y = (mean),
    fill = SampleType)) + geom_col() + geom_text(data = subset(fungi_order_comp_2,
    mean_label > 1), aes(y = lab_ypos, label = paste(mean_label,
    "%", sep = ""), group = SampleType), fontface = "bold", size = 6,
    color = "white")

p + theme(axis.text.x = element_text(angle = -70, hjust = 0,
    vjust = 0.5)) + theme(text = element_text(size = 34)) + ylab("Mean Relative Abundance") +
    guides(fill = guide_legend(title = "Sample Type")) + scale_fill_viridis_d(option = "C",
    begin = 0.85, end = 0) + xlab("Order") + scale_x_discrete(labels = c(o__Capnodiales = "Capnodiales"
    o__Eurotiales = "Eurotiales", o__Glomerellales = "Glomerellales",
    o__Hypocreales = "Hypocreales", o__Pleosporales = "Pleosporales",
```

```
    o__Filobasidiales = "Filobasidiales", o__Sporidiobolales = "Sporidiobolales",
    o__Dothideales = "Dothideales", o__Mucorales = "Mucorales"))
```

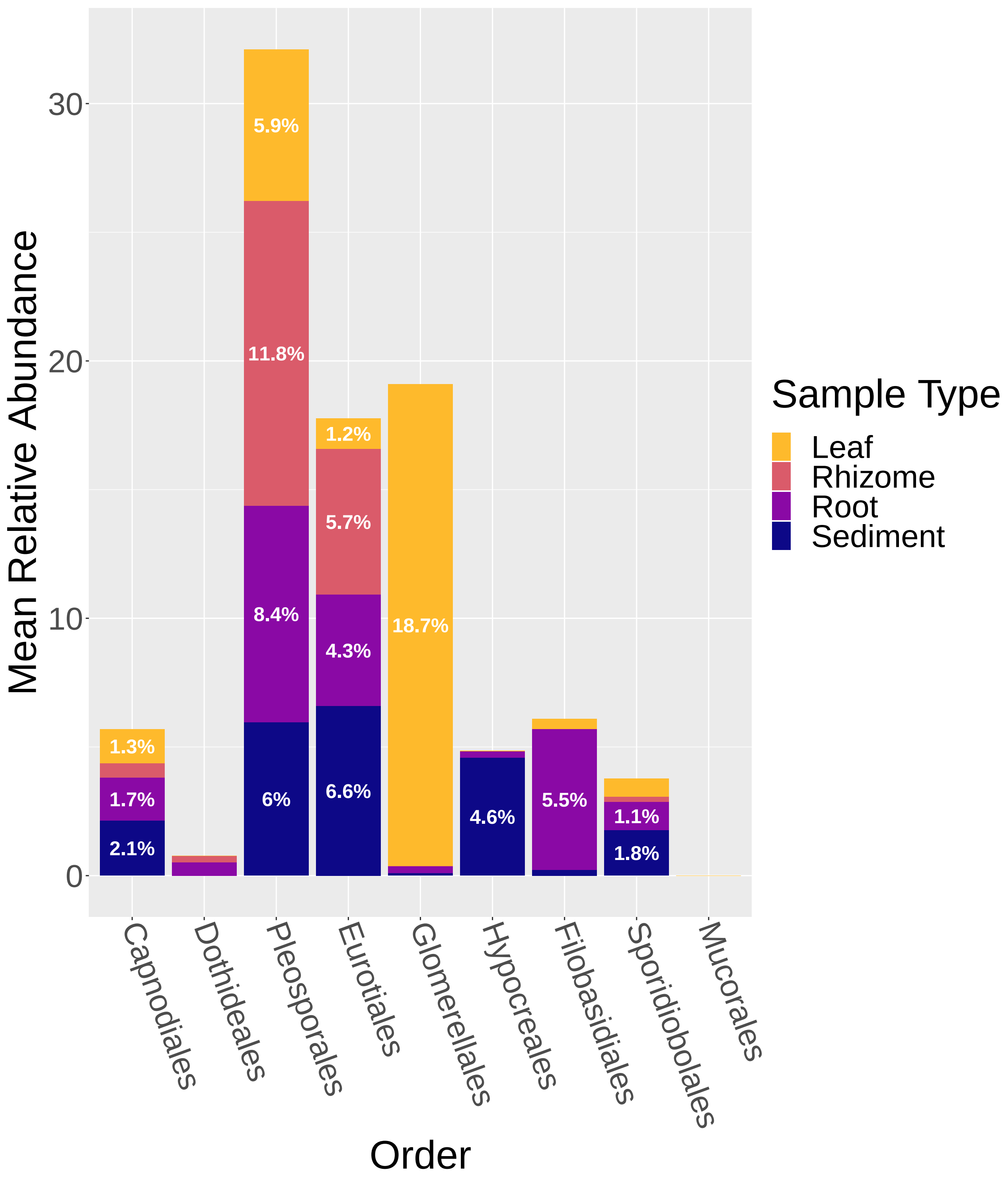


```
# ggsave(filename = 'Fun_Hist_ITS.pdf', plot = last_plot(),
# device = 'pdf', width = 12, height = 16, dpi = 300)
```

#### Comparing the prescence / abscence of the fungal genera isolated from each sample type in this study to the ITS amplicon data

```
# collapse the ASV table to genus level
SGFungi_rare10000_taxglom_GenusLevel_RA <- tax_glom(SGFungi_rare10000_RA,
    taxrank = "Genus", NArm = FALSE)

# Extract only genera of interest (those isolated here)
IsolateMIOnly <- subset_taxa(SGFungi_rare10000_taxglom_GenusLevel_RA,
    Genus == "g__Cladosporium" | Genus == "g__Mycosphaerella" |
        Genus == "g__Aureobasidium" | Genus == "g__Penicillium" |
        Genus == "g__Talaromyces" | Genus == "g__Rhodotorula" |
        Genus == "g__Colletotrichum" | Genus == "g__Acrostalagmus" |
        Genus == "g__Emericellopsis" | Genus == "g__Sarocladium" |
        Genus == "g__Trichoderma" | Genus == "g__Naganishia" |
        Genus == "g__Pseudozyma" | Genus == "g__Absidia")

# Convert the phyloseq object to an R dataframe
IsolateMIOnly_df <- psmelt(IsolateMIOnly)

# Summarize the df and calculate P/A for each sample type
grouped_genus <- group_by(IsolateMIOnly_df, SampleType, Genus)
avgs_genus <- summarise(grouped_genus, PA_ITS = 1 * (sum(Abundance) >
    0))

# rename the genera from the ASV table to match the isolate
# file
avgs_genus <- avgs_genus %>% mutate(Genus = recode(Genus, g__Cladosporium = "Cladosporium sp",
    g__Mycosphaerella = "Ramularia sp", g__Aureobasidium = "Aureobasidium sp",
    g__Penicillium = "Penicillium sp", g__Talaromyces = "Talaromyces sp",
    g__Rhodotorula = "Rhodotorula sp", g__Acrostalagmus = "Acrostalagmus sp",
    g__Colletotrichum = "Colletotrichum sp", g__Emericellopsis = "Emericellopsis sp",
    g__Sarocladium = "Sarocladium sp", g__Trichoderma = "Trichoderma sp",
    g__Naganishia = "Naganishia sp"))


# Read in csv containing isolate information (same as used
# earlier)
fungi_genus <- read.csv("fungi_tab1.csv")

# Subset to only include isolates able to be identified to
# the genus level Removes Hypocreales sp and Pleosporales sp
fungi_only_genus <- subset(fungi_genus, Molecular.ID == "Cladosporium sp" |
    Molecular.ID == "Ramularia sp" | Molecular.ID == "Aureobasidium sp" |
    Molecular.ID == "Penicillium sp" | Molecular.ID == "Talaromyces sp" |
    Molecular.ID == "Rhodotorula sp" | Molecular.ID == "Colletotrichum sp" |
    Molecular.ID == "Acrostalagmus sp" | Molecular.ID == "Emericellopsis sp" |
    Molecular.ID == "Sarocladium sp" | Molecular.ID == "Trichoderma sp" |
    Molecular.ID == "Naganishia sp")


# Remove sample types not represented in ITS data
fungi_only_genus <- filter(fungi_only_genus, Tissue != "Seawater")
```

```r
fungi_only_genus <- filter(fungi_only_genus, Tissue != "Leaves and roots")
fungi_only_genus <- filter(fungi_only_genus, Tissue != "Roots and rhizome")

# Rename Molecular.ID to Genus
fungi_only_genus$Genus <- fungi_only_genus$Molecular.ID

# Rename Tissue to SampleType
fungi_only_genus$SampleType <- fungi_only_genus$Tissue

# Count number of isolates for each sample type
fungi_count <- fungi_only_genus %>% count(Genus, SampleType)

# Summarize the df and calculate P/A for each sample type
grouped_isolate <- group_by(fungi_count, SampleType, Genus)

avgs_isolate <- summarise(grouped_isolate, PA_Isolate = 1 * (sum(n) >
    0))


# Combine the two P/A results (the isolate data and the ITS
# amplicon data)
combined_data <- full_join(avgs_isolate, avgs_genus, by = c("Genus",
    "SampleType"))
```

```
## Warning: Column `Genus` joining factors with different levels, coercing to
## character vector

## Warning: Column `SampleType` joining factors with different levels, coercing to
## character vector
```

```r
# Replace NAs with 0s
combined_data <- combined_data %>% mutate(PA_ITS = replace_na(PA_ITS,
    0))
combined_data <- combined_data %>% mutate(PA_Isolate = replace_na(PA_Isolate,
    0))

# sum, 0 = not detected on that tissue, 1 = found in either
# ITS or isolate data but not both, 2 = found in both
# datasets
combined_data$PA_Overall <- combined_data$PA_Isolate + combined_data$PA_ITS

# Reorder fungal genera to be alphabetical by phylum / class
# / order
combined_data$Genus_f = factor(combined_data$Genus, levels = c("Cladosporium sp",
    "Ramularia sp", "Aureobasidium sp", "Penicillium sp", "Talaromyces sp",
    "Rhodotorula sp", "Acrostalagmus sp", "Colletotrichum sp",
    "Emericellopsis sp", "Sarocladium sp", "Trichoderma sp",
    "Naganishia sp"))

# Reorder sample types to match histograms
combined_data$SampleType = factor(combined_data$SampleType, levels = c("Leaf",
    "Rhizome", "Root", "Sediment"))

# Make heatmap
s <- ggplot(combined_data, aes(SampleType, Genus_f, fill = factor(PA_Overall))) +
```

```
    geom_tile() + scale_fill_manual(values = c("grey90", "grey50",
    "black"), labels = c("Not detected", "One method", "Both methods"),
    name = "Detection Level")
s <- s + theme(axis.text.x = element_text(angle = -70, hjust = 0,
    vjust = 0.5)) + theme(text = element_text(size = 34))
s + scale_y_discrete(labels = c(`Cladosporium sp` = "Cladosporium",
    `Ramularia sp` = "Ramularia", `Aureobasidium sp` = "Aureobasidium",
    `Penicillium sp` = "Penicillium", `Talaromyces sp` = "Talaromyces",
    `Rhodotorula sp` = "Rhodotorula", `Acrostalagmus sp` = "Acrostalagmus",
    `Colletotrichum sp` = "Colletotrichum", `Emericellopsis sp` = "Emericellopsis",
    `Sarocladium sp` = "Sarocladium", `Trichoderma sp` = "Trichoderma",
    `Naganishia sp` = "Naganishia")) + xlab("Sample Type") +
    ylab("Genus")
```

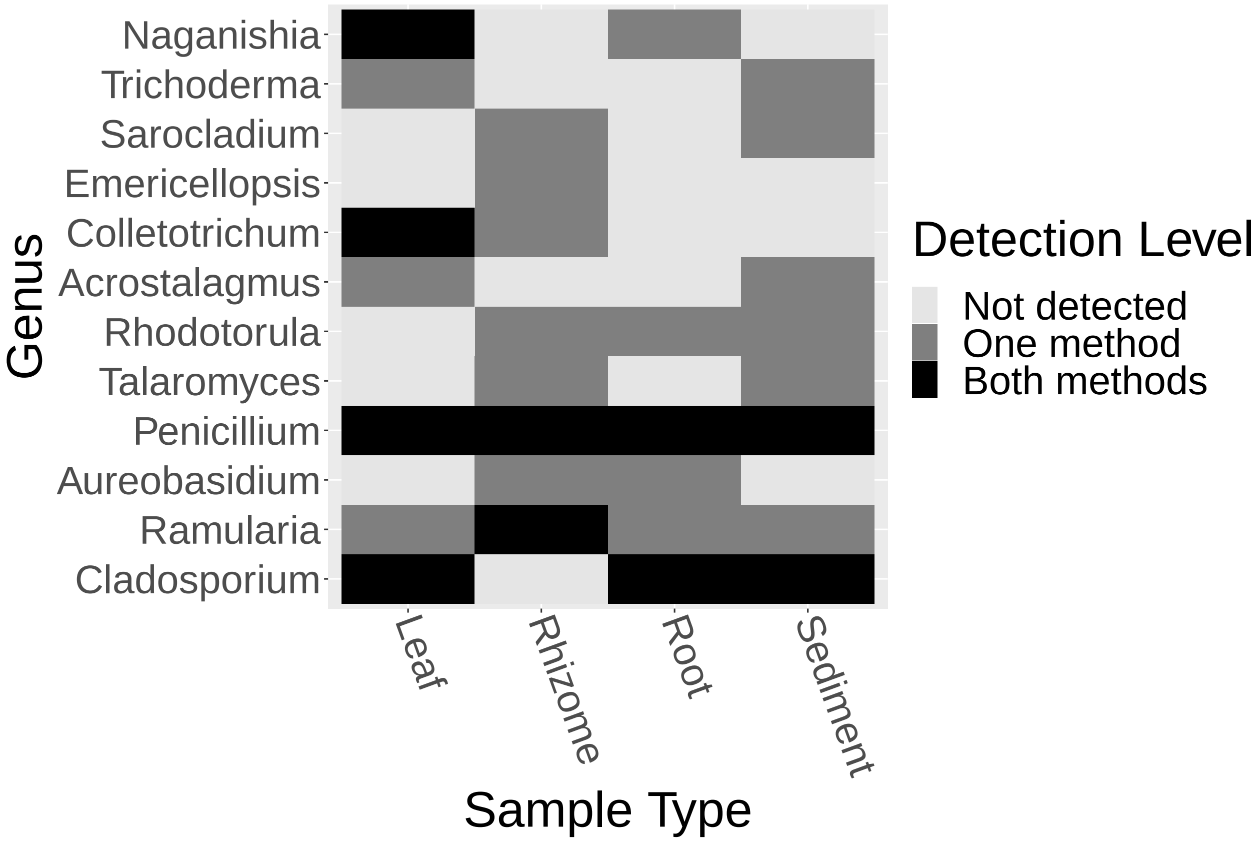


```
# ggsave(filename = 'Fun_Heatmap.pdf', plot = last_plot(),
# device = 'pdf', width = 12, height = 8, dpi = 300)
```
